# Synthetic multistability in mammalian cells

**DOI:** 10.1101/2021.02.10.430659

**Authors:** Ronghui Zhu, Jesus M. del Rio-Salgado, Jordi Garcia-Ojalvo, Michael B. Elowitz

## Abstract

In multicellular organisms, gene regulatory circuits generate thousands of molecularly distinct, mitotically heritable states, through the property of multistability. Designing synthetic multistable circuits would provide insight into natural cell fate control circuit architectures and allow engineering of multicellular programs that require interactions among cells in distinct states. Here we introduce MultiFate, a naturally-inspired, synthetic circuit that supports long-term, controllable, and expandable multistability in mammalian cells. MultiFate uses engineered zinc finger transcription factors that transcriptionally self-activate as homodimers and mutually inhibit one another through heterodimerization. Using model-based design, we engineered MultiFate circuits that generate up to seven states, each stable for at least 18 days. MultiFate permits controlled state-switching and modulation of state stability through external inputs, and can be easily expanded with additional transcription factors. Together, these results provide a foundation for engineering multicellular behaviors in mammalian cells.

## Main Text

Multistability is one of the hallmarks of multicellular life, allowing genetically identical cells to exist in thousands of molecularly distinct and mitotically stable cell types or states (*1*–*4*). Understanding natural multistable circuits and engineering synthetic ones have been longstanding challenges in developmental and synthetic biology (*5*–*22*). While key regulatory interactions have been identified in some natural fate control circuits, it generally remains unclear what minimal circuitry is sufficient to generate the observed multistability. On the synthetic side, engineered multistable circuits would provide a foundation for synthetic developmental and therapeutic circuits that could generate a spectrum of designed cell types with specialized functions. However, previous efforts in mammalian cells have been limited to two-state systems and used architectures that cannot be easily expanded to larger numbers of states (*8*, *9*, *12*). To address both issues, we sought to engineer a synthetic multistable circuit in mammalian cells based on prevalent features of natural fate control systems.

An ideal synthetic multistable system would allow cells to remain in any of a set of distinct expression states over extended timescales of many cell cycles, despite biological noise. In addition, it would provide three key capabilities exhibited by its natural counterparts (Fig. 1A): First, it would permit transient external inputs to switch cells between states, similar to the way signaling pathways direct fate decisions during development (*23*). Second, it would support control over the number and stability of different states, and enable irreversible transitions, similar to those that occur during natural differentiation (*20*, *24*). Third, it would be expandable (scalable), so that one could increase the number of states by introducing additional components without re-engineering an existing functional circuit, analogous to expansions in the number of cell types and states that have occurred during evolution (*25*, *26*).

**Fig. 1.**
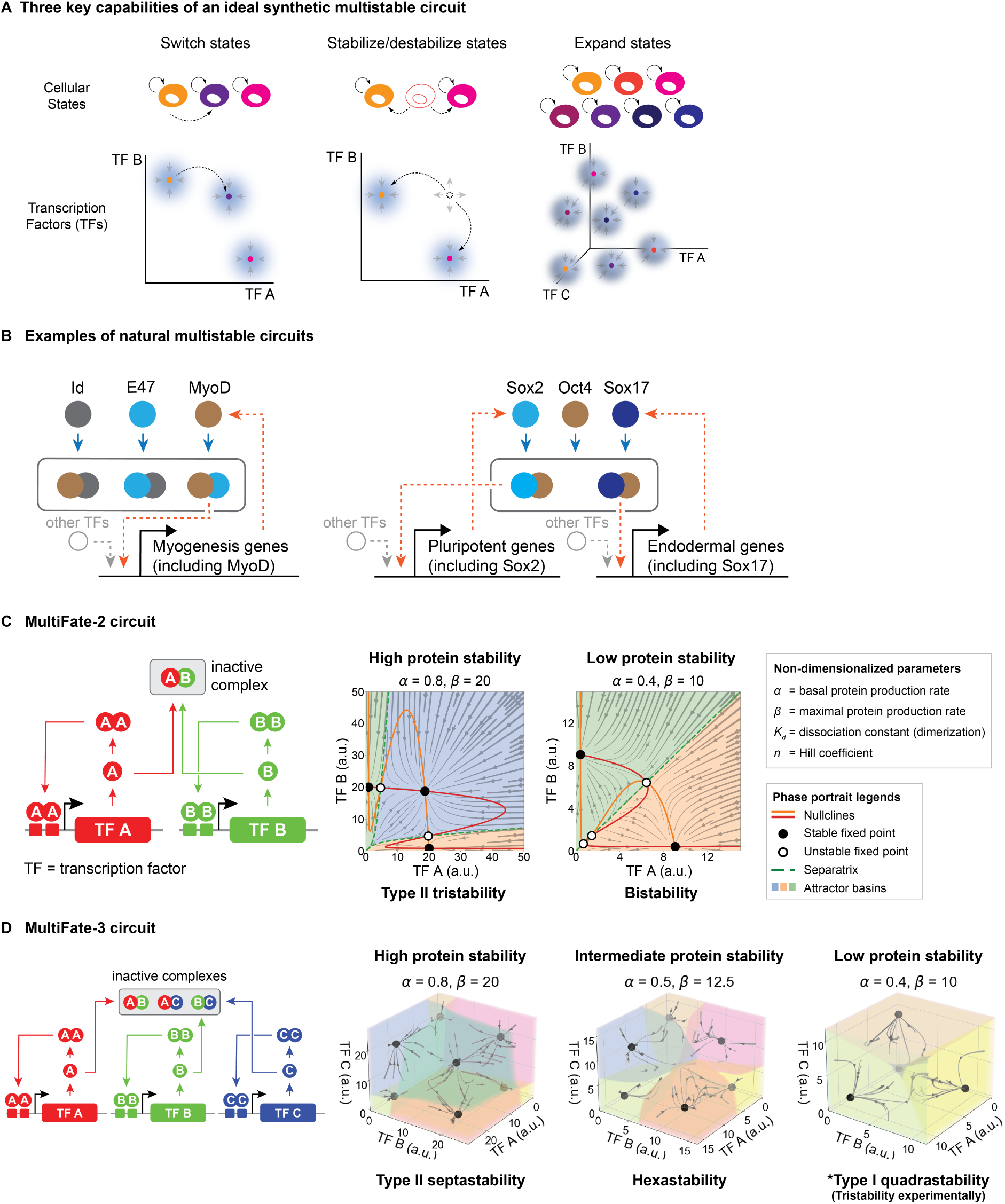
The naturally-inspired MultiFate architecture generates diverse types of multistability in the model. (A) An ideal synthetic multistable circuit should generate multiple stable states, represented by colored cell cartoons (upper level) and attractors in the phase space of transcription factor expression (lower level, schematic, TF A-C on coordinate axes represent transcription factor concentrations); support control of state-switching (left); and state stability (middle); and allow easy expansion of states by addition of more transcription factors (right). (B) Competitive protein-protein interactions and autoregulatory feedback are prevalent in natural multistable circuits that control myogenesis (left) and endodermal differentiation (right). These simplified and abridged diagrams highlight the role of transcriptional autoregulation and promiscuous dimerization in these circuits. Blue arrows indicate competitive protein-protein interactions, which can involve higher order multimerization. Orange dashed arrows indicate direct or indirect positive transcriptional feedback. (C, D) Models of the MultiFate-2 circuit and MultiFate-3 circuit (Box 1, Supplementary Materials) generates diverse types of multistability in different parameter regimes (indicated above plots). In the model of the MultiFate-3 circuit, reduced protein stability generates 4 stable states (type I quadrastability), but the state in which all transcription factors are OFF has a very small attractor basin, and thus should be effectively unstable in the presence of biological noise. Therefore only 3 states should be experimentally stable in a low protein stability regime, consistent with experimental results in Fig. 4B, Low TMP columns. Complete lists of multistability regimes are shown in Fig. S1 and S2. For both panels, each axis represents the total concentration of each transcription factor. All models used here are non-dimensionalized, with *K*_*d*_ = 1 and *n* = 1.5. Note that in the non-dimensionalized model, changing protein stability is equivalent to multiplying and with the same factor (Box 1).

Natural mammalian multistable circuits provide inspiration for such a synthetic architecture (Fig. 1B) (*27*–*37*). For example, during myogenesis, muscle regulatory factors such as MyoD heterodimerize with E proteins to activate their own expression and the broader myogenesis program, while Id family proteins heterodimerize with muscle regulatory factors or E proteins to disrupt this process (*34*–*36*). Similarly, during embryogenesis, Sox2 and Sox17 competitively interact with the pluripotency regulator Oct4 to control the early differentiation decision between pluripotency and endodermal differentiation (*31*, *32*). These and other circuits thus appear to use transcription factors that autoregulate their own expression and cross-regulate each other’s expression, and competitively interact to form a variety of homodimers, heterodimers, and higher order multimeric forms. Related combinations of positive autoregulation and cross-inhibition could extend multistability behaviors beyond bistability and generate bifurcation dynamics that explain the partial irreversibility of cell differentiation (*11*, *16*, *38*–*41*). Nevertheless, it remains unclear whether these natural architectures could be adapted to enable synthetic multistability. Here, we show how a synthetic multistable system based on principles derived from natural cell fate control systems can generate robust, controllable, expandable multistability in mammalian cells.

## Results

### The MultiFate circuit architecture generates diverse types of multistability through a set of promiscuously interacting, autoregulatory dimer-dependent transcription factors

Inspired by natural fate control circuits, we designed the MultiFate system, in which transcription factors homodimerize to positively autoregulate their own expression, and heterodimerize to mutually inhibit each other’s transcriptional activity (Fig. 1C). The transcription factors share a common dimerization domain, allowing them to competitively form both homodimers and heterodimers. Further, the promoter of each transcription factor gene contains binding sites that can be strongly bound only by the homodimeric form of its own protein, allowing homodimer-dependent self-activation. By contrast, heterodimers do not efficiently bind to any promoter in this design. Heterodimerization thus acts to mutually inhibit the activity of both constituent transcription factors.

Mathematical modeling shows how the MultiFate architecture provides each of the desired capabilities described above (Fig. 1A) in physiologically reasonable parameter regimes (Box 1, Table S4, Supplementary Materials). A MultiFate circuit with just two transcription factors, designated MultiFate-2, can produce diverse types of multistability containing 2, 3 or 4 stable fixed points depending on protein stability and other parameter values (Fig. 1C, Fig. S1A). In particular, a regime designated type II tristability, that includes stable states expressing either A, B, or both, is analogous to multilineage priming in uncommitted progenitor cells, with the double positive state playing the role of a multipotent progenitor (*42*–*50*). In the model, transient expression of one transcription factor can switch cells between states (Fig. S3, Movie S1). Reducing the protein stability of transcription factors can cause bifurcations that selectively destabilize specific states (Fig. 1C). Finally, the model is expandable: addition of a new transcription factor to the MultiFate-2 model generates a three transcription factor “MultiFate-3” circuit that supports additional stable states with the same parameter values (Fig. 1D, Fig. S2A). Together, these modeling results suggest that the MultiFate architecture can support a rich array of multistable behaviors.

### Engineered zinc fingers enable homodimer-dependent activation and heterodimer-dependent inhibition

Synthetic zinc finger (ZF) transcription factors provide an ideal platform to implement the MultiFate circuit. Zinc finger DNA-binding domains can be assembled from individual zinc fingers to recognize target DNA binding sites with high specificity (*51*). When fused with a transcriptional activation domain, the resulting zinc finger transcription factors can activate gene expression when bound to its promoter (*52*–*55*). Further, engineered ZF DNA-binding domain containing three fingers bind weakly as monomers to 9bp target sites, but can bind much more strongly as homodimers to 18bp tandem binding site pairs (*56*). This property allows homodimer-dependent transcriptional activity and potentially allows inhibition through heterodimerization.

To engineer ZF transcription factors, we started with ZF-GCN4-AD, a synthetic transcription factor containing the ErbB2 ZF DNA-binding domain fused to a GCN4 homodimerization domain and a VP48 transcriptional activation domain (Fig. 2A) (*56*). As a monomeric (non-dimerizing) control, we also constructed a variant of this protein lacking GCN4, termed ZF-AD. To assay their transcriptional activity, we constructed a reporter containing two repeats of 18bp tandem binding site pairs driving the expression of a Citrine fluorescent protein (*56*). We then co-transfected each of the transcription factors together with the reporter and an mTagBFP2 co-transfection marker into CHO-K1 cells, and analyzed Citrine expression by flow cytometry 36 hours later (Fig. 2A, Fig. S4A, Supplementary Materials). The Wild-type (WT) ZF-GCN4-AD factors strongly activated the reporter, as desired, while the monomeric variant ZF-AD exhibited weaker, but still undesirable, basal activity (Fig. 2A, Fig. S4B). Following early structural work and recent advances in zinc finger engineering (*52*, *57*–*59*), we incorporated arginine-to-alanine mutations at key positions in the ZF known to impact DNA binding, which decreased background monomeric activity without reducing foreground homodimer activity (bars within red square in Fig. 2A, Fig. S4B). Further, we replaced the GCN4 with the FKBP12F36V (FKBP) homodimerization domain (*60*). The resulting FKBP-ZF-AD factor allows dose-dependent control of dimerization with the activating drug AP1903 (Fig. 2B). Finally, using this general design, we constructed eight additional dimer-dependent ZF transcription factors, and validated orthogonal DNA-binding specificities for four of them (Fig. S4C). Together, these results provided a set of orthogonal, externally controllable, homodimer-dependent ZF transcription factors.

**Fig. 2.**
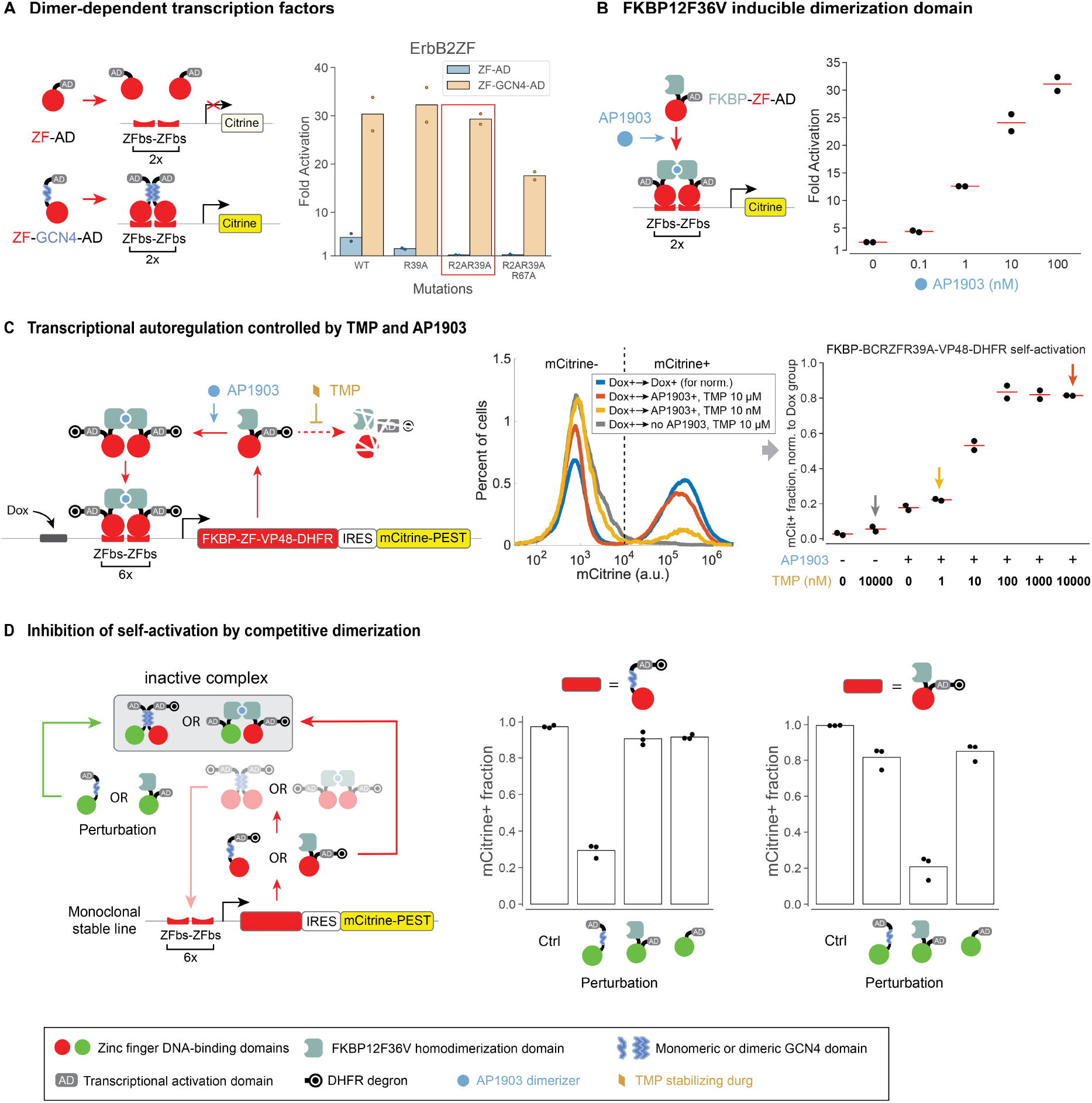
Engineered transcription factors enable homodimer-dependent autoregulation and heterodimerization-based mutual inhibition. (A) Zinc finger transcription factors can generate homodimer-dependent activation. (Left) Schematic representations of test constructs, in which ZF DNA-binding domains (ErbB2ZF (*56*), red circle) fused to activation (VP48, AD) and in some cases dimerization (GCN4, blue squiggle) domains bind to corresponding target sites (red pads) to activate reporter expression (Citrine). Activators (Table S1) were expressed from a constitutive CAG promoter (*87*), and co-transfected with a Citrine fluorescence reporter that has 2 repeats of 18bp tandem binding site pairs at the promoter. (Right) Arginine-to-alanine mutations in the ErbB2ZF domain modulated reporter activation by ZF-GCN4-AD and ZF-AD. The R2AR39A variant was selected due to high ZF-GCN4-AD activation and minimal ZF-AD activation. Fold activation is defined in Fig. S4A. WT = wild-type variant. (B) Using FKBP12F36V (FKBP) as the homodimerization domain (light cyan partial box), activation can be controlled by AP1903 (blue circle) in a dose-dependent manner. Orange circle = BCRZFR39A. (C) Transcription factor self-activation can be controlled by TMP and AP1903. (Left) Design of the controllable self-activation circuit (Table S1). (Right) Stable polyclonal cells showed bimodal mCitrine distribution upon circuit activation. An empirical threshold at mCitrine=10^4^ separates the population into mCitrine- and mCitrine+ subpopulations, and the mCitrine+ fraction normalized to constitutive Dox treatment group (Dox+, gray curve) was used to quantify self-activation strength (Supplementary Materials). Colored arrows indicate data from the middle panel. IRES = internal ribosome entry site; PEST = constitutive degradation tag; (D) Self-activation was inhibited by competing transcription factors with a different ZF and matching dimerization domains. Two monoclonal stable lines could spontaneously turn on self-activation in media containing AP1903 and TMP (Table S2). Perturbation is introduced into monoclonal lines by stably integrating plasmids expressing different transcription factor variants into monoclonal lines (Supplementary Materials). Red circle = 42ZFR2AR39AR67A; Green circle = BCRZFR39A. In all panels, each dot represents one replicate, and each red line or bar indicates the mean of replicates.

The MultiFate circuit design requires that each transcription factor positively autoregulates its own expression in a homodimer-dependent manner. To validate this capability, we designed a self-activation construct (Fig. 2C, left), in which a transcription factor with FKBP dimerization domain is expressed from a promoter containing multiple repeats of its own 18bp homodimeric binding site (Table S1). This construct allowed independent Dox-inducible activation through upstream Tet3G (Takara Bio) binding sites. It also incorporated a dihydrofolate reductase (DHFR) degron (*61*), which can be inhibited by the drug trimethoprim (TMP), at the C-terminus of transcription factor, permitting control of protein stability. Thus, it allowed independent control of dimerization, expression, and degradation. To enable dynamic single-cell readout of expression, we also incorporated a destabilized mCitrine fluorescent protein on the same construct. Finally, we integrated this construct into Tet3G-expressing CHO-K1 cells, generating a stable polyclonal population for further analysis (Table S2, Supplementary Materials).

To test for positive autoregulation, we transiently induced transcription factor expression for 24 hours with Dox, and then withdrew Dox and checked whether cells could sustain circuit activation, at different dimerization strengths and protein stability, controlled by AP1903 and TMP, respectively. In the presence, but not the absence, of AP1903, cells exhibited a bimodal distribution of mCitrine fluorescence, with well-separated peaks (Fig. 2C, middle and right), consistent with dimer-dependent positive autoregulation in a subset of cells. TMP, by stabilizing transcription factors, also promoted self-activation in a dose-dependent manner (Fig. 2C, Fig. S5A). Together, these results show that a single self-activating dimer-dependent transcription factor can generate bimodal dynamics in a controllable manner.

MultiFate’s final requirement is the ability of one transcription factor to effectively inhibit another through formation of inactive heterodimers. To test this, we selected monoclonal cell lines expressing the self-activating circuit, and then stably integrated constructs expressing proteins with a different ZF DNA-binding domain and a matching or mismatching dimerization domain (Table S1, Table S2, Supplementary Materials). Only proteins with matching dimerization domains inhibited the self-activating transcription factor, consistent with inhibition through heterodimerization (Fig. 2D, Fig. S5B). Taken together, these results provided a set of engineered ZF transcription factors that exhibit homodimer-dependent activation and heterodimeric inhibition with external control of dimerization and protein stability.

### The MultiFate-2 circuit generates tristability

To construct a complete MultiFate circuit, we selected two dimer-dependent transcription factors, henceforth designated A and B, with distinct DNA binding specificities but the same FKBP homodimerization domain (Table S1, Supplementary Materials), and expressed them from promoters containing multiple repeats of their corresponding 18bp homodimeric binding sites. The promoters also incorporated Tet3G or ERT2-Gal4 response elements (*62*) to allow independent external induction. A and B were co-expressed with destabilized mCherry or mCitrine fluorescent proteins, respectively, for visualization (Fig. 3A). We stably integrated both genes simultaneously in CHO-K1 cells expressing Tet3G and ERT2-Gal4 proteins, and further characterized three stable monoclonal cell lines, designated MultiFate-2.1, MultiFate-2.2 and MultiFate-2.3 with different promoter configurations (Table S2, Supplementary Materials).

**Fig. 3.**
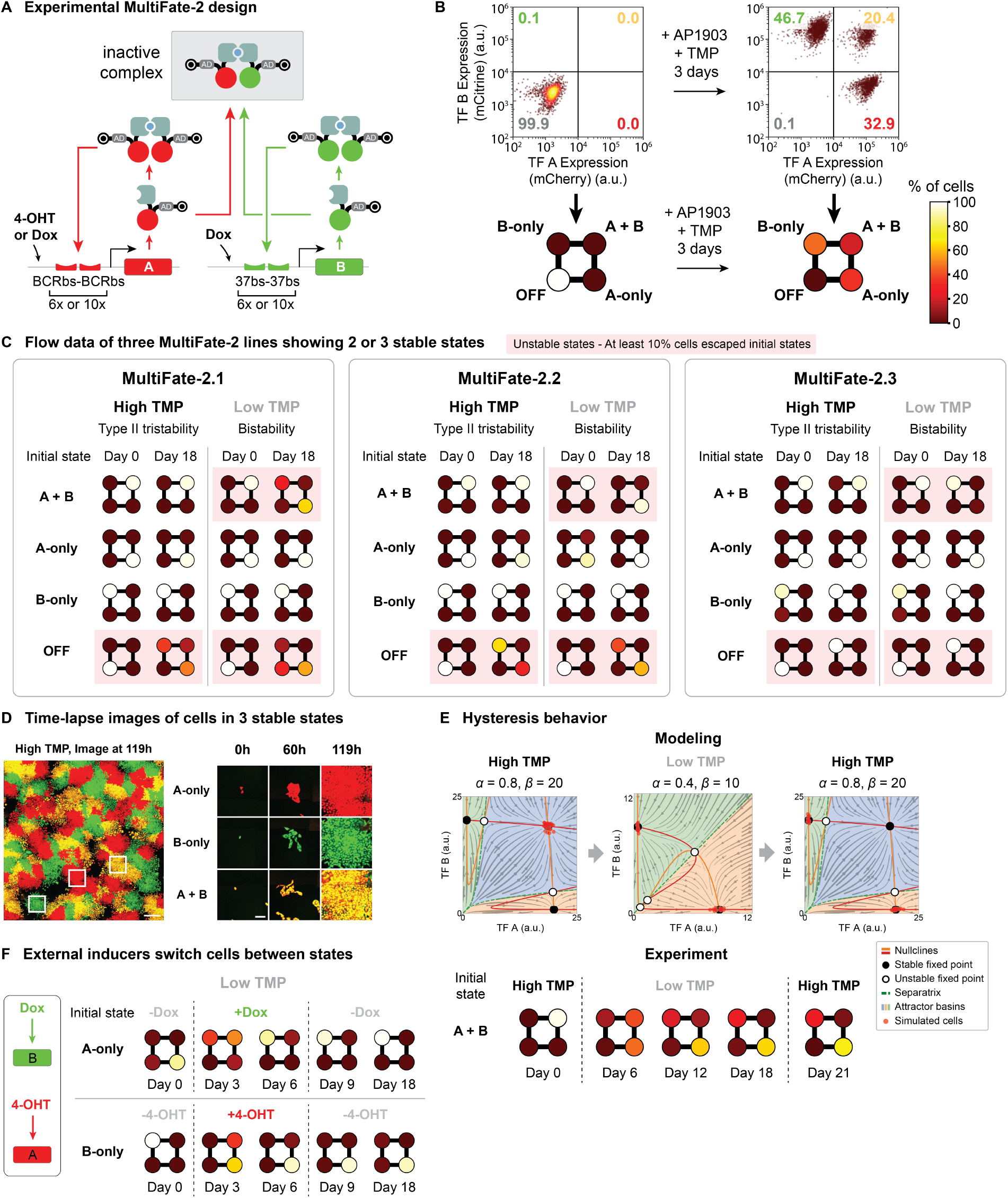
Experimental data show that MultiFate-2 generates multiple stable states, supports modulation of state stability and allows state-switching. (A) The experimental MultiFate-2 design uses two self-activation cassettes differ only in their ZF DNA-binding domains and binding sites, and fluorescent proteins. Each cassette expresses FKBP-ZF-VP16-DHFR-IRES-FP-PEST, where ZF represents either BCRZFR39A or 37ZFR2AR11AR39AR67A and FP represents either mCherry or mCitrine, for A and B, respectively. Detailed construct maps and differences among MultiFate-2 lines are available in Table S1 and Table S2. (B) MultiFate-2.1 cells spontaneously activate A, B or both cassettes upon addition of 100 nM AP1903 and 10 μM TMP. Cell percentages in OFF, A-only, B-only and A+B states were quantified and plotted as a square with four colored circles representing the percentage of cells in each quadrant (Supplementary Materials). (C) Three MultiFate-2 lines exhibited type II tristability in the High TMP condition, and bistability in the Low TMP condition. In all conditions, AP1903 concentration is 100 nM. Exact concentrations of TMP are shown in Fig. S6-8. Unstable states, defined by states having more than 10% cells escaping after 18 days, were marked in pink rectangles. (D) A-only, B-only and A+B states were each stable during growth from single MultiFate-2.3 cells into colonies over 5 days under a time-lapse microscope. Scale bar: 500 μm for the wide field image (left), 100 μm for zoomed in images (right). (E) Escape from the destabilized A+B state was irreversible, as shown by both modeling (Movie S3), and experiment using MultiFate-2.1 cells. (Top) The model used here is non-dimensionalized, with *K*_*d*_ = 1 and *n* = 1.5. Simulated cells on phase portraits were calculated using the Gillespie stochastic simulation algorithm (*89*) (Supplementary Materials). (Bottom) The Day 0 and Day 18 squares are the same with those in panel C (MultiFate-2.1, A+B initial state under Low TMP), since data are from the same experiment. (F) MultiFate-2.3 cells can be switched between states by transient 4-OHT or Dox treatment. Cells were cultured in 100 nM AP1903 + 30 nM TMP (Low TMP condition) throughout the experiment. 4-OHT = 25 nM, Dox = 500 ng/ml. In all panels, initial A-only, B-only and A+B cells were sorted from a population of cells in different states, while initial OFF cells came from cells in regular CHO media without any inducers. Each square represents the mean fractions of three replicates.

We next sought to test whether MultiFate circuits support multistability in general and, more specifically, whether they could operate in the type II tristability regime, which exhibits features resembling multilineage priming and permits irreversible bifurcations to bistability regime to generate hysteresis (*42*–*50*) (Box 1). We activated the circuit by transferring MultiFate-2.1 cells to media containing AP1903 and TMP, allowing dimerization and stabilizing the transcription factors. As expected in the regime of type II tristability (Box 1, Fig. 1C), cells went from low expression of both transcription factors (OFF state) to one of three distinct states, with either A, B, or both transcription factors highly expressed (Fig. 3B), which we designated as A-only, B-only and A+B states, respectively. These states were well separated, by ~25-50 fold differences in either mCherry or mCitrine expression. To assess their stability, we sorted cells from each of these states and cultured them continuously for 18 days (Supplementary Materials). Strikingly, nearly all cells remained in the sorted state for this extended period (Fig. 3C, Fig. S6). Stability required the positive autoregulatory circuit, as withdrawal of AP1903 and TMP collapsed expression of both factors within 2 days (Fig. S6). Similar overall behavior was also observed in MultiFate-2.2 and MultiFate-2.3 (Fig. 3C, Fig. S7 and Fig. S8A). All three MultiFate-2 cell lines thus exhibited dynamics strikingly consistent with type II tristable behavior (Fig. 1C).

Time-lapse imaging provided a more direct view of multistability. We cultured single cells from different initial states in the same well and imaged them as they developed into colonies (Fig. 3D). In almost all colonies (132/134), all cells maintained their initial states for the full duration of the movie, at least 5 days or 7-8 cell cycles (Fig. 3D, Fig. S9, Movie S2). Together with the flow cytometry analysis, these results demonstrate that all three MultiFate-2 lines can sustain long-term tristability.

### MultiFate-2 supports modulation of state stability and allows controlled state-switching

In natural cell fate control systems, destabilization of multipotent states has been suggested to facilitate irreversible transitions into differentiated fates (*16*, *41*). Similarly, in the MultiFate-2 model, reducing protein stability specifically destabilizes the A+B state, shifting the system from type II tristability to bistability (Fig. 1C). To test whether this predicted bifurcation would occur in the experimental system, we transferred A-only, B-only and A+B cells from media containing high TMP concentrations (“High TMP”) to similar media with reduced TMP concentrations (“Low TMP”), which decreases protein stability by permitting degron function.

As predicted, reducing protein stability selectively destabilized the A+B state, but not the A-only and B-only states (Fig. 3C, Low TMP columns), causing cells to continuously transit from the destabilized A+B state to the stable A-only or B-only states (Fig. S6, Bottom). Different MultiFate-2 cell lines exhibited different transition biases, reflecting clone-specific asymmetries in the experimental MultiFate-2 systems, in a manner consistent with an asymmetric MultiFate model (Movie S3, Supplementary Materials). Also as predicted, the OFF state remained unstable in the Low TMP condition, although cells exited the unstable OFF state more slowly than they did under the High TMP condition (Fig. 3C, OFF row). Reduced protein stability thus generated the predicted tristable-to-bistable bifurcation.

The ability of a transient stimulus to trigger an irreversible fate change is a hallmark of many cell fate control systems, including oocyte maturation (*20*), and exit from pluripotency (*24*). In the model, transiently reducing protein stability produces an irreversible (hysteretic) response: cells initially in the A+B state transit to A-only or B-only states when protein stability is reduced, and remain there, rather than return to the A+B state, when protein stability is restored to its initial value. Stochastic simulations of single cell dynamics exhibit this hysteretic behavior (Fig. 3E, top; Movie S3). To test whether hysteretic dynamics occur in the experimental system, we analyzed cell populations in the MultiFate-2.1 line after reducing TMP concentration and then restoring it to its initial high concentration. Indeed, escape from the destabilized A+B state was irreversible, as cells remained in the A-only or B-only state even after they were transferred back to the High TMP condition (Fig. 3E, bottom; Fig. S6). MultiFate’s ability to support irreversible transitions allows it to produce behaviors resembling stem cell differentiation.

Finally, we asked to what extent we could deliberately switch cells from one state to another through transient perturbations. In MultiFate-2.3, the A and B genes are independently inducible by 4-hydroxy-tamoxifen (4-OHT) and Dox (Table S2). Transient Dox treatment transitioned A-only cells to the B-only state within 6 days (Fig. 3F, top). During switching, cells briefly expressed both proteins at elevated levels (Fig. S8B), consistent with model predictions (Fig. S3, Movie S1). Conversely, transient 4-OHT induction of A expression transitioned B-only cells to the A-only state in a similar manner (Fig. 3F, bottom). MultiFate-2 thus allows controlled state-switching with transient inputs. Taken together, these results demonstrate that MultiFate-2 circuits allow modulation of state stability, irreversible (hysteretic) cell state transitions, and direct control of state-switching with external inputs.

### MultiFate is expandable

Because MultiFate circuits implement mutual inhibition among transcription factors through heterodimerization, MultiFate circuits can be expanded simply by adding additional transcription factors, without re-engineering existing components. In the model, adding a third transcription factor to a MultiFate-2 circuit produces a range of new stability regimes containing 3, 4, 6, 7, or 8 stable fixed points, depending on parameter values (Fig. 1D, Fig. S2, Movie S4, Supplementary Materials).

Can experimental MultiFate-2 circuits be similarly expanded? To answer this question, we stably integrated a third ZF transcription factor, denoted C, containing the same FKBP dimerization domain as A and B, co-expressed with a third fluorescent protein, mTurqoise2, into the MultiFate-2.2 cell line (Fig. 4A, Table S2, Supplementary Materials). After addition of AP1903 and TMP, the resulting MultiFate-3 cells populated 7 distinct clusters, termed A-only, B-only, C-only, A+B, A+C, B+C, and A+B+C states (Fig. 4B). Most cells went to the B-only state (79.5%±0.3%), reflecting asymmetries within the circuit (e.g. higher basal expression at B cassettes). The set of stable states, and the absence of a stable OFF state with no proteins expressed, strikingly resembled the type II septastable regime at high protein stability (Fig. 1D, Fig. S2A). To assess the stability of these states, we sorted cells from each of the seven states, and continuously cultured them in media containing AP1903 and TMP, analyzing the culture every 3 days by flow cytometry. Remarkably, each of the seven states was stable for the full 18-day duration of the experiment (Fig. 4B, High TMP columns; Fig. S10). Long-term stability required AP1903 and TMP, as expected (Fig. S13). Finally, cells from each state could be reset by withdrawal of AP1903 and TMP and then re-differentiated into all of the states when AP1903 and TMP were added back (Fig. S13). This indicates that the observed stability is not the result of a mixture of clones permanently locked into distinct expression states.

**Fig. 4.**
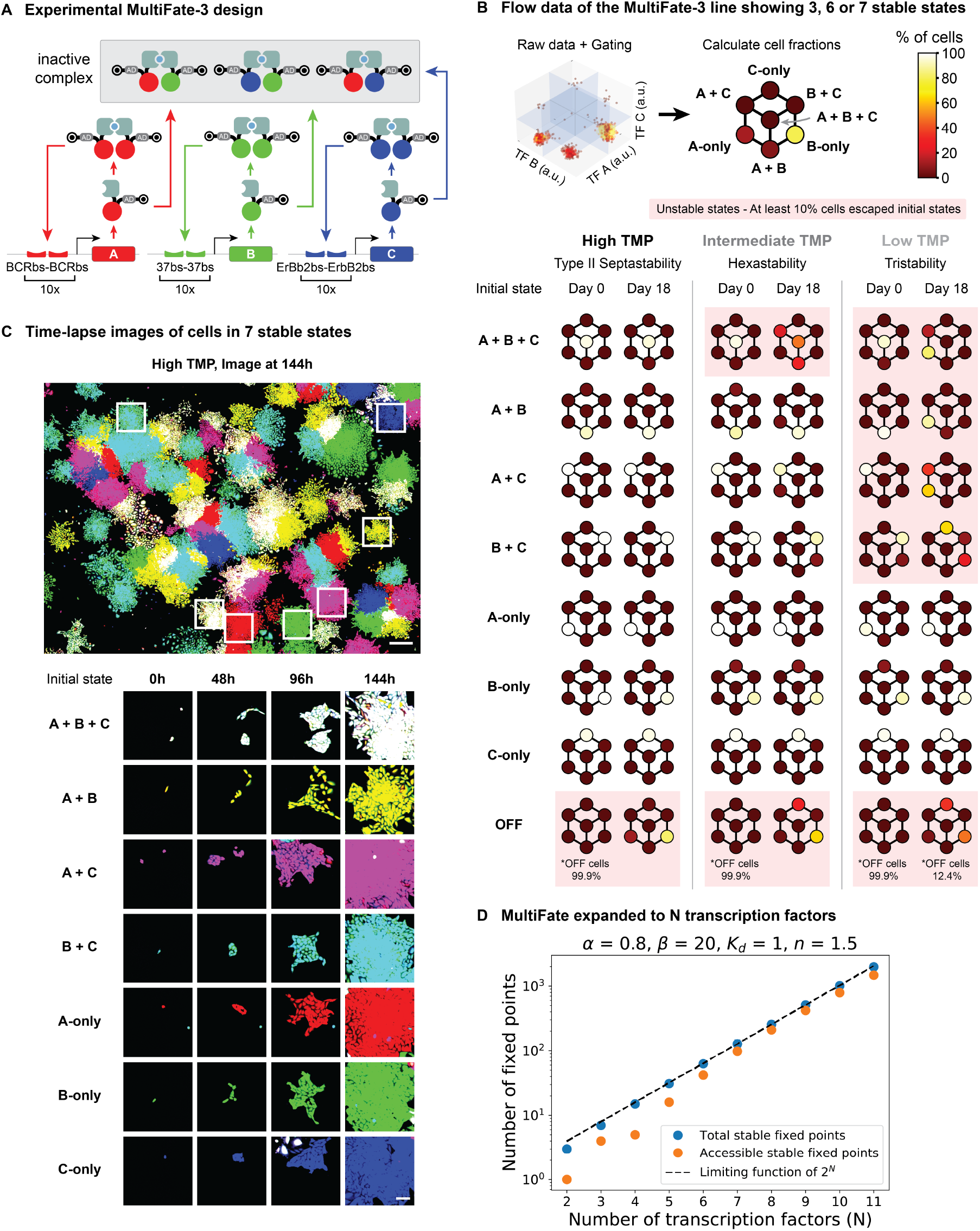
MultiFate architecture is expandable to include three and potentially even more transcription factors. (A) The experimental MultiFate-3 design uses three self-activation cassettes differ only in their ZF DNA-binding domains and binding sites, and fluorescent proteins. Each cassette expresses FKBP-ZF-VP16-DHFR-IRES-FP-PEST, where ZF represents either BCRZFR39A, 37ZFR2AR11AR39AR67A or ErbB2ZFR2AR39A, and FP represents either mCherry, mCitrine or mTurquoise2, for A, B and C, respectively. Detailed construct maps are available in Table S1. (B) The MultiFate-3 line exhibited type II septastability, hexastability and tristability in three different TMP conditions. (Top) State percentages except OFF state percentage were then quantified and plotted as a hexagon with seven colored circles representing the percentage of cells in each octant (Supplementary Materials). OFF state percentages were usually low (<1%) across all conditions, and are separately labeled only when their fractions exceed 1%. (Bottom) High TMP condition = 100 nM AP1903 + 100 nM TMP; Intermediate TMP condition = 100 nM AP1903 + 40 nM TMP; Low TMP condition = 100 nM AP1903 + 10 nM TMP. Except for OFF state cells, cells in different initial states were sorted from a mixed population of cells in the High TMP condition. Initial OFF cells came from cells in regular CHO media without any inducers. Circle colors in each hexagon represent the mean percentages of three replicates. (C) Cells in each of the seven states were stable during growth from single cells into colonies over 6 days under a time-lapse microscope. Scale bar: 500 μm for the wide field image (left), 100 μm for zoomed in images (right). (D) MultiFate can be expanded (model). The number of accessible stable fixed points grows monotonically with the number, N, of transcription factors species in the model. An accessible stable fixed point is defined as a stable fixed point that has an attractor basin volume that is at least 25% of maximal attractor basin volume of stable fixed points in the same system (same N). The parameter set provided above the plot is the same non-dimensionalized parameter set used in MultiFate-2 and MultiFate-3 models under high protein stability.

To directly visualize the septastable dynamics of MultiFate-3, we co-cultured single cells sorted from each of the seven states and performed live imaging as they grew into colonies. Consistent with the flow cytometry results, cells retained their initial states for the full 6-day duration of the experiment in almost every colony (153/157) (Fig. 4C, Fig. S9, Movie S5).

Like MultiFate-2, the number and stability of different states in MultiFate-3 can be modulated. In the model, reducing protein stability repeatedly bifurcates the system from type II septastability (7 stable states) through hexastability (6 stable states) to tristability (3 stable states) (Fig. 1D). This process resembles the progressive loss of cell fate potential during stem cell differentiation (*63*– *66*).

To experimentally test this prediction, we transferred cells in each of the 7 states cultured under the High (100 nM) TMP (high protein stability) condition to similar media with Intermediate (40 nM) or Low (10 nM) TMP conditions. As predicted by the model, the Intermediate TMP condition destabilized only the A+B+C state, but not the other 6 states (Fig. 4B, Intermediate TMP columns, Fig. S11), while the Low TMP condition destabilized all multi-protein states, preserving only those in which a single transcription factor is expressed (Fig. 4B, Low TMP columns, Fig. S12A). Consistent with the model, these transitions were also irreversible: restoring High TMP concentrations did not cause cells to repopulate previously destabilized states (Fig. S14). Taken together, these results demonstrate that the MultiFate-3 circuit supports septastability, and allows controlled bifurcations to produce irreversible cell state transitions.

Can the MultiFate architecture be expanded beyond three transcription factors? In principle, with larger numbers of transcription factors, accumulation of basal expression could limit the ability of any transcription factor species to reach sufficiently high concentrations. Additionally, even if more stable states are generated, many could have relatively small attractor basins, rendering them effectively inaccessible. To find out how higher order MultiFate systems behave, we modeled circuits containing up to 11 transcription factor species, using the same parameter values established for MultiFate-2 and MultiFate-3. Even in larger MultiFate circuits, basal expression of more transcription factors did not curtail multistability. In fact, the number of stable fixed points grew nearly exponentially with the number of transcription factors, N, approaching a limiting function of 2N (Fig. 4D). To assess the accessibility of these states, we quantified their attractor basin volumes, defining states as “accessible” when their attractor basins were at least 25% as large as the maximal basin of other fixed points in the same system. Remarkably, most of the stable fixed points were accessible. For instance, with N=11 transcription factors, 1474 of the 1981 stable fixed points were accessible. More generally, the number of accessible fixed points grew monotonically with N, at a rate approaching that of the total number of fixed points (Fig. 4D, Fig. S15). These results indicate that the MultiFate architecture can be expanded to generate large numbers of accessible stable states.

## Discussion

The astonishing diversity of cell types in our own bodies underscores the critical importance of multistable circuits and provokes the fundamental question of how to engineer a robust, controllable, and expandable synthetic multistable system. Here, we took inspiration from two ubiquitous features of natural multistable systems, namely competitive protein-protein interactions and transcriptional autoregulation, to design a synthetic multistable architecture that operates in mammalian cells. The MultiFate circuits introduced here exhibit many of the hallmarks of natural cell fate control systems. They generate as many as seven molecularly distinct, mitotically heritable cell states from three transcription factors (Fig. 3 and 4). They allow controlled switching of cells between states with transient transcription factor expression (Fig. 3F), similar to fate reprogramming (*67*). They support modulation of state stability (Fig. 3 and 4) and permit irreversible (hysteretic) cellular transitions through externally controllable parameters such as protein stability (Fig. 3E, Fig. S14), similar to the irreversible loss of cell fate potential during stem cell differentiation (*68*, *69*). Finally, implementing cross-inhibition at the protein levels makes MultiFate expandable, allowing one to exponentially increase the number of states in the system by ‘plugging in’ additional transcription factors, without re-engineering the existing circuit, a useful feature for synthetic biology. The same design principle may play a related role in natural systems, allowing the emergence of new cell states through transcription factor duplication and subfunctionalization in a manner analogous to the stepwise expansion of MultiFate circuits demonstrated here (*32*, *70*–*76*).

A remarkable feature of this circuit is its close agreement with predictions from a dynamical systems model (Box 1). Despite a lack of precise quantitative parameter values for many molecular interactions, the qualitative behaviors possible with this circuit design can be enumerated and explained from simple properties of the components and their interactions. More precise measurements of effective biochemical parameters and stochastic fluctuations could help to explain, eliminate, or exploit asymmetries (Fig. 3C and Fig. 4B) and provide a better understanding of the timescales of state transitions.

MultiFate has a relatively simple structure, requiring a modest number of genes, all of the same type, yet exhibits robust memory behaviors, scalability, and predictive design. Future work should extend MultiFate into a full-fledged synthetic cell fate control system, in which extrinsic signals can be used to navigate cells sequentially through a series of fate choices, recapitulating cell behaviors associated with normal development. Coupling MultiFate to synthetic signaling systems such as synNotch (*77*, *78*), MESA (*79*), synthekines (*80*), orthoIL-2/2R (*81*), engineered GFP (*82*) and auxin (*83*) should enable flexible and orthogonal control. MultiFate could also allow engineering of multicellular cell therapeutic programs. For example, one could engineer a stem-like state that can either self-renew or “differentiate” into other states that recognize and remember different input signals and communicate with one another via signaling to coordinate complex response programs. Such strategies will benefit from the ability of MultiFate to allow probabilistic differentiation into multiple different states in the same condition (Fig. 3C, 4B). In this way, we anticipate that the MultiFate architecture will provide a scalable foundation for exploring the circuit-level principles of cell fate control and enabling new multicellular applications in synthetic biology.

#### Box 1. Design of the MultiFate circuit

Here we introduce the mathematical model of the MultiFate circuit and show how it can be used to design the experimental system and predict its behavior. For simplicity, we focus on a symmetric MultiFate-2 circuit whose two transcription factors share identical biochemical parameters and differ only in their DNA binding site specificity. A similar analysis of systems with more transcription factors and asymmetric parameters is presented in Supplementary Materials.

We represent the dynamics of protein production and degradation using ordinary differential equations (ODEs) for the total concentrations of the transcription factors A and B, denoted [*A*_*tot*_] and [*B*_*tot*_], respectively. We assume that the rate of production of each protein follows a Hill function of the corresponding homodimer concentration, [*A*_2_] or [*B*_2_], with maximal rate β, Hill coefficient *n*, and half-maximal activation at a homodimer concentration of *K*_*M*_. A low basal rate of “leaky” protein production, denoted α, is included to allow self-activation from low initial expression states. Finally, each protein can degrade and dilute (due to cell division) at a total rate δ, regardless of its dimerization state. To simplify analysis, we non-dimensionalize the model by rescaling time in units of δ^−1^, and concentrations in units of *K*_*M*_ (Supplementary Materials). We can then write:

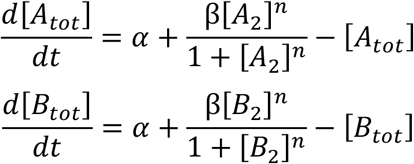

Dimerization dynamics occur on a faster timescale than protein production and degradation (*84*). This separation of timescales permits us to assume that the distribution of monomer and dimer states remains close to their equilibrium values, generating the following relationships between the concentrations of monomers, [*A*] and [*B*], and dimers, [*A*_2_], [*B*_2_], and [*AB*]:

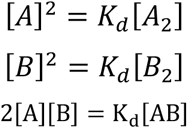

Here, because the two transcription factors share the same dimerization domain, homo- and hetero-dimerization are assumed to occur with equal dissociation constants, *K*_*d*_. Additionally, conservation of mass implies that [*A*_*tot*_] = [*A*] + [*AB*] + 2[*A*_2_], with a similar relationship for B. Introducing the equilibrium equations given above into this conservation law produces expressions for the concentrations of the activating homodimers in terms of the total concentrations of A and B (which do not change form after non-dimensionalization):

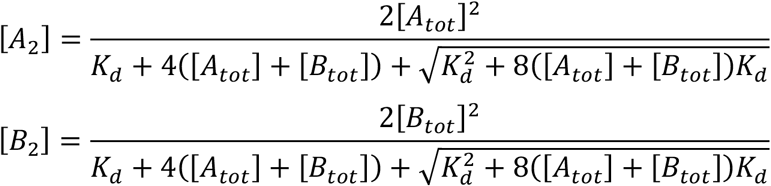

Inserting these expressions into the differential equations for [*A*_*tot*_] and [*B*_*tot*_] above, we obtain a pair of coupled ordinary differential equations with only [*A*_*tot*_] and [*B*_*tot*_] as variables.

To understand the behavior of this system in physiologically reasonable parameter regimes (Table S4, Supplementary Materials), we used standard approaches from dynamical systems analysis (*85*). We plotted the nullclines (Fig. 1C, solid lines), defined by setting each of the ODEs above to zero. We then identified fixed points at nullcline intersections, and determined their linear stability (Fig. 1C, black and white dots) (*85*). Finally, we delineated the basins of attraction for each stable fixed point (Fig. 1C, shaded regions).

Using this analysis, we identified parameter values that support type II tristability, a regime that minimally embodies the developmental concept of multilineage priming (*42*–*50*), and permits transitions to a bistable regime in response to tuning of a single parameter, thus allowing one to explore bifurcation dynamics (Fig. 1C, Fig. S1B). Stronger self-activation (higher values of β) was more likely to produce type II tristability (Fig. S1B, β row and column). Too much leaky production (high α) allowed both transcription factors to self-activate, reducing the degree of multistability, while too little (low α) stabilized the undesired OFF state (Fig. S1B, α column). Strong dimerization (low *K*_*d*_) was essential for type II tristability (Fig. S1B, *K*_*d*_ row and column). Finally, a broad range of Hill coefficients *n* ≥ 1 were compatible with type II tristability. While higher values of *n* led to a reduced sensitivity to other parameters and allowed the system to tolerate higher values of α, they also stabilized the OFF state (Fig. S1B, *n* row and column). Together, these results suggested that an ideal design would maximize β, minimize *K*_*d*_, and use intermediate values of α and *n*.

Based on these conclusions, we incorporated multiple repeats of the homodimeric binding sites to maximize β, used strongly associating FKBP12F36V homodimerization domains (*60*) to minimize *K*_*d*_, and modified the promoter sequences to allow some leaky expression to optimize α (Supplementary Materials). Finally, while we did not directly control *n*, we expected that the repeated homodimeric binding sites should lead to modest ultrasensitivity (*86*). These design choices produced the selected type II tristability in the experimental system (Fig. 3C, Supplementary Materials).

A key feature of the MultiFate design is its ability to qualitatively change its multistability properties through bifurcations in response to parameter changes. In particular, the mathematical model predicts that protein stability can control the number of stable fixed points in phase space. In the non-dimensionalized model, the protein degradation rate, δ, does not appear explicitly but enters through the rescaling of α and β by (δ*K*_*M*_)^−1^ (see “non-dimensionalization of MultiFate model” section in Supplementary Materials). Thus, tuning protein stability is equivalent to multiplying both α and β by a common factor, which we term the “protein stability factor”. Reducing protein stability shifts the nullclines closer to the origin, causing the two unstable fixed points to collide with the stable A+B fixed point in a subcritical pitchfork bifurcation (Fig. 1 and Fig. S1B, “protein stability factor” column) (*87*). The result is a bistable system with A-only and B-only stable fixed points at somewhat lower concentrations (Fig. 1C). To experimentally realize this bifurcation, we designed the circuit to allow external control of transcription factor protein stability using the drug-inducible DHFR degron (Fig. 2C). As predicted, reducing protein stability destabilized the A+B state, but preserved the A-only and B-only stable states.

In this way, model-based design enabled us to rationally engineer tristability as well as externally controllable transitions to bistability in the experimental system (Fig. 3C, Fig. 3E). The model presented above also generalizes in a straightforward manner to allow analysis of expanded MultiFate circuits (Fig. 1D, Supplementary Materials).

