## Supplementary Materials for "Synthetic multistability in mammalian cells"

#### Materials and Methods

Plasmid construction

Constructs used in this study are listed in Table S1. Some constructs were generated using standard cloning procedures. The inserts were generated using PCR or gBlock synthesis (IDT) and were annealed by Gibson assembly with backbones that are linearized using restriction digestion. The rest of the constructs were designed by the authors and synthesized by GenScript. All the construct maps are available at data.caltech.edu/records/1882, and we've initiated the process to deposit selected constructs used to build MultiFate lines onto Addgene.

Tissue culture

CHO-K1 cells were cultured at 37$^oC$ in a humidity controlled chamber with 5% $CO_2$. The growth media consisted of Alpha MEM Earle's Salts (FujiFilm Irvine Scientific) supplemented with 10% FBS, 1 U/ml penicillin, 1 μg/ml streptomycin and 1 mM L-glutamine. For experiments requiring a change of inducer conditions, cells were first washed 3 times using media with the new inducer condition. After the wash, cells were rinsed once with Dulbecco's Phosphate-Buffered Saline (DPBS, Life Technologies) and trypsinized with 0.25% Trypsin (Life Technologies) for 3 min at 37$^oC$. Trypsinized cells were then transferred into a new well with media added with the new inducer condition.

Transient transfection

24 hours before transfection, $0.05x10^6$ CHO-K1 cells were seeded per well of a 24-well plate using standard culture media. The next day, cells were transfected with plasmids using Lipofectamine LTX and PLUS Reagents (Thermo fisher) according to manufacturer's protocol.

Cell line construction

Stable cell lines used in this study are listed in Table S2**.** Stable cell lines were generated using the PiggyBac Transposon system (System Biosciences, (*89*)). CHO-K1 cells in a 24-well plate were co-transfected with transgene constructs in a PiggyBac expression backbone, an EF1α-PuroR plasmid and a Super PiggyBac Transposase plasmid (System Biosciences). Cells were transferred into a 6-well plate and selected with 10 μg/ml puromycin for 3 days to obtain a stable polyclonal population. To make monoclonal lines, polyclonal cells were either counted and diluted to roughly 1 cell per well, or sorted by FACS as single cells into 384-well plates. In either case, the plates were checked under microscope after 4-5 days to eliminate wells without cells or with more than one colony. For wells that only have a single colony growing, cells were expanded and subsequent screening was performed to obtain multistable clones.

To identify potential MultiFate-2 clones that could operate in several multistability regimes, we turned to our MultiFate mathematical model for intuition. Through parameter screening we found that when one progressively reduces protein stability starting from a value at which the state with all transcription factors expressed simultaneously (all-ON) was stable could generate a progressive reduction of multistability (Fig. S1B).

To achieve similar behavior experimentally, we sought to identify MultiFate-2 monoclones that exhibit stable A+B state. To test whether the A+B state was stable in different monoclones, we first transiently induce the expression of all transcription factors by Dox and 4-OHT for 12 hours, then washed and replaced with media containing 100 nM AP1903 and 10 μM (saturating concentration) TMP. After 3 days, we checked (by mCherry and mCitrine expression) whether cells could sustain the expression of both transcription factors under a fluorescence microscope. Monoclones that could maintain the stability of A+B state were selected as MultiFate-2 clones for further analysis. Using a similar method, a monoclonal that could maintain the stability of the A+B+C state was selected as the MultiFate-3 line. All the MultiFate lines used in the current study are available from the corresponding author.

Flow cytometry

All samples were harvested from 24-well plates. Cells were first rinsed with 500 μl DPBS, and then trypsinized with 75 μl 0.25% trypsin for 3 min at 37$^{o}$C. Trypsin was neutralized by resuspending cells in 300 μl flow cytometry buffer containing Hank's Balanced Salt Solution (Life Technology) and 2.5 mg/ml Bovine Serum Albumin. Cell samples were then filtered by 40 μm cell strainers and analyzed by a flow cytometer (CytoFLEX, Beckman Coulter). We used the EasyFlow Matlab-based software package developed by Yaron Antebi to process flow cytometry data (https://antebilab.github.io/easyflow/).

Characterization of ZF transcription factors

To characterize transcriptional activation of different ZF transcription factor variants (Fig. 2A, Fig 2B and Fig. S4), we transfected CHO-K1 cells in 24-well plates with 50ng ZF transcription factor plasmid (Table S1, construct MF08–MF62), 50ng reporter plasmid (Table S1, construct MF01–MF07) and 25ng EF1α-mTagBFP2. In the "ReporterOnly" group, ZF transcription factor plasmid was replaced by an empty plasmid with only a constitutive promoter but no ZF transcription factor. In the "NoReporter" well, both ZF transcription factor plasmid and reporter plasmid were replaced by an empty plasmid. For ZF transcription factors with FKBP homodimerization domain (Fig. 2B), AP1903 was added to the transfection media. 36 hours after transfection, cells were harvested and analyzed by flow cytometry. To maximize the reporter dynamic range, we selected and compared highly transfected cells by gating cells with high levels of a BFP co-transfection marker. Median citrine fluorescence intensity of gated cells was used to calculate fold activation. To calculate fold activation, median fluorescence values of NoReporter samples, representing the cellular autofluorescence background, were first subtracted from ReporterOnly and Reporter+ZF samples. The ratio between background-subtracted Reporter+ZF value and background-subtracted ReporterOnly value is then the fold activation of that ZF transcription factor on that reporter.

Characterization of ZF transcription factor self-activation

Each self-activation construct (Table S1, construct MF63–MF69) was stably integrated into Tet3G-expressing CHO-K1 cells. After puromycin selection, polyclonal cells were transferred into media containing 500 ng/ml Dox to transiently express ZF transcription factors. After 24 hours of Dox treatment, cells were washed 3 times with regular media and transferred into media containing different concentrations of AP1903 and/or TMP to test how dimerization and/or protein stability affect self-activation. One sample of cells (Dox+ sample) continued to be cultured in 500 ng/ml Dox as the positive control. After another 72 hours, cells were harvested and analyzed by flow cytometry. Stable polyclonal cells showed strongly bimodal mCitrine distribution upon circuit activation (Fig. 2C, middle). We used an empirical threshold at mCitrine=$10^4$ fluorescence units to separate the population into mCitrine- (cells with no circuit integrated or integrated circuit cannot self-activate) and mCitrine+ (cells with integrated circuit activated) subpopulations. To only consider cells with at least one stably integrated activatable circuit, we normalized the mCitrine+ fraction of each sample to the mCitrine fraction of Dox+ sample, in which high concentrations of Dox should turn on all stably integrated activatable circuits. This normalized mCitrine+ fraction was used to compare self-activation strength across different AP1903 and TMP combinations.

<u>Assay showing inhibition of self-activation by competitive dimerization</u>

Two monoclonal self-activation stable lines (with 42ZFR2AR39AR67A DNA-binding domain and either GCN4 or FKBP as the homodimerization domains, see Table S2) were selected since they showed spontaneous and homogeneous self-activation upon the addition of 100 nM AP1903 and 10 μM TMP (Fig. S5B). To whether competitive dimerization inhibits self-activation (Fig. 2D and Fig. S5B), we stably integrated plasmids (Table S1, construct MF72–MF80) expressing different transcription factor variants and a co-translational mCherry in these two monoclonal lines. After puromycin selection, cells were transferred into media containing 100 nM AP1903 + 10 μM TMP to permit self-activation, and measured by flow cytometry after another 72 hours. The mCherry+ cell population was gated for analysis. We introduced protein variants through stable integration, instead of transient transfection, to avoid nonspecific transcriptional interference by transient high expression of proteins during transfection, and to test inhibition of self-activation in a cellular environment better mimicking the MultiFate-2 and MultiFate-3 stable cell lines.

<u>Fluorescence Activated Cell Sorting (FACS)</u>

To separate MultiFate cells in distinct states for subsequent experiments (Fig. 3 and Fig. 4), cells were harvested and resuspended in sorting buffer (BD FACS Pre-Sort Buffer) supplemented with 1 U/ml DNAse I, AP1903 and TMP. Cells were then sorted into media containing different concentrations AP1903 and TMP, according to the experiments. Cell sorting was performed by Caltech Flow Cytometry Facility.

<u>Flow cytometry measurement of long-term state stability of MultiFate cells</u>

To characterize long-term state stability of MultiFate cells in different media conditions (Fig. 3 and Fig. 4), we trypsinized and transferred 4% of cells into fresh media with the same condition every three days. The remaining cells were resuspended in flow cytometry buffer and analyzed by a flow cytometer. The resulting two-dimensional (or three-dimensional for MultiFate-3) fluorescence intensity plots were then divided into four quadrants (or eight octants) by an empirical threshold in each of the two (or three) fluorescence channels. The exact values of empirical thresholds for different MultiFate cell lines are slightly different due to expression differences and are shown in the figure legends of Fig. S6-8 and Fig. S10. The percentage of cells in quadrants (or octants) were then calculated for each sample. The mean percentage of cells across three samples was plotted as a square (or a hexagon) with colored circles representing the percentages. For the MultiFate-3 line, cell percentages in OFF state were usually very low and were separately labeled if the percentage was greater than 1%.

<u>Time-lapse imaging</u>

To visualize state stability of MultiFate cells (Movie S2 and Movie S5), MultiFate cells were sparsely seeded in an imaging 24-well plate (μ-Plate 24 Well Black, ibidi) with regular media. After 6 or 12 hours, the media was replaced with media containing AP1903 and TMP. Time-lapse images were acquired on an inverted Olympus IX81 fluorescence microscope with Zero Drift Control (ZDC), an ASI 2000XY automated stage, a Photometrics 95B camera (Teledyne Photometrics) and a 20x UPlanS/Apo objective (0.75 NA, Olympus). Fluorescent proteins were excited with an X-Cite XLED1 light source (Lumen Dynamics). Images were automatically acquired every one hour, controlled by MetaMorph software (Molecular Devices). Cells were kept in a custom-made environmental chamber enclosing the microscope, controlling a humidified, 37°C and 5% $CO_2$ atmosphere. Media was changed every three days.

Attractor basin analysis of MultiFate circuit with N transcription factors

In theory, the attractor basin volume of a stable fixed point goes to infinity (except for the fixed point with all transcription factors OFF) if the concentration of each transcription factor has no limit. However, transcription factor concentrations are bounded by their maximum expression levels. Consequently, attractor basin volumes are finite and volumes of different basins can be compared. In the non-dimensionalized model, transcription factor concentrations are confined to the interval $[\alpha, \alpha + \beta]$, which corresponds to the equilibrium concentrations when a transcription factor is not self-activating and when it is fully self-activating, respectively. To calculate the approximate volumes of attractor basins, we initialized an $N$-dimensional concentration grid with $k$ points in each dimension, spaced at equal linear intervals, for a total of $k^N$ points. Using these grid points as initial conditions, we numerically solved the differential equations to compute forward trajectories to their final stable fixed point, using the expanded MultiFate-N model (see below). We labeled each grid point based on which stable fixed point its trajectory ends at. The attractor basin volume of a stable fixed point was then approximated as

$$V_i = \frac{\text{number of grid points that end at fixed point i}}{k^N} \times \beta^N$$

where $\beta^N$ represents the total phase space volume.

Parameter screening of MultiFate-2 and MultiFate-3 circuits

Each parameter dependency plot in Fig. S1B and Fig. S2B represents a field of 100×100 points. The color of each point denotes the multistability type generated through MultiFate-2 (Fig. S1B) or MultiFate-3 (Fig. S2B) with the indicated parameter combination. To identify the multistability type of each parameter set, we initialized a 2- or 3-dimensional grid, for MultiFate-2 and MultiFate-3, respectively, with eight values for each transcription factor concentration. Using these grid points as starting points, we csomputed trajectories using the MultiFate differential equations. We then grouped the end points of these trajectories into clusters and used the centers (center of mass) of these clusters as estimated locations of stable fixed points. We double-checked the stability of the estimated fixed points using standard linear stability analysis (*85*) type by the locations of these stable fixed points.

Data and code availability

All data generated and all the computational and data analysis and modeling code used in the current study are available at data.caltech.edu/records/1882.

**Supplementary Text**

Non-dimensionalization of MultiFate model

As shown in Box 1, each ordinary differential equation (ODE) for $[A_{tot}]$ and $[B_{tot}]$ consists of three terms: (i) a basal protein production rate α, (ii) a Hill function describing self-activation dynamics with maximal rate β, Hill coefficient $n$, and half-maximal activation at a homodimer concentration of $K_M$, and (iii) a protein degrade rate δ. We can then write:

$$\frac{d[A_{tot}]}{dt} = \alpha + \frac{\beta[A_2]^n}{K_M^n + [A_2]^n} - \delta[A_{tot}]$$

$$\frac{d[B_{tot}]}{dt} = \alpha + \frac{\beta[B_2]^n}{K_M^n + [B_2]^n} - \delta[B_{tot}]$$

Dimerization dynamics occur on a faster timescale than protein production and degradation (*84*). This separation of timescales permits us to assume that the distribution of monomer and dimer states remains close to equilibrium, generating the following relationships between the concentrations of monomers ($[A]$ and $[B]$), and dimers ($[A_2]$, $[B_2]$, and $[AB]$) based on the law of mass action:

$$[A]^2 = K_d[A_2]$$

$$[B]^2 = K_d[B_2]$$

$$2[A][B] = K_d[AB]$$

Here, because the two transcription factors share the same dimerization domain, homo- and hetero-dimerization are assumed to occur with equal dissociation constants, $K_d$. When deriving these three equations from the law of mass action, each monomer is counted twice in homodimerization reactions, and is counted once in the heterodimerization reaction, thus a factor of two is introduced in the third equation to account for this statistical difference. Additionally, conservation of mass implies that $[A_{tot}] = [A] + [AB] + 2[A_2]$, with a similar relationship for B.

Solving these equations produces expressions for the concentrations of the activating homodimers in terms of the total concentrations of A and B:

$$[A_2] = \frac{2[A_{tot}]^2}{K_d + 4([A_{tot}] + [B_{tot}]) + \sqrt{K_d^2 + 8([A_{tot}] + [B_{tot}])K_d}}$$

$$[B_2] = \frac{2[B_{tot}]^2}{K_d + 4([A_{tot}] + [B_{tot}]) + \sqrt{K_d^2 + 8([A_{tot}] + [B_{tot}])K_d}}$$

To non-dimensionalize the model, we rescale time in units of $\delta^{-1}$, and concentrations in units of $K_M$. Then we have:

$$t \leftarrow t\delta, \qquad [A_{tot}] \leftarrow [A_{tot}]/K_M, \qquad [B_{tot}] \leftarrow [B_{tot}]/K_M, \qquad [A_2] \leftarrow [A_2]/K_M,$$

$$[B_2] \leftarrow [B_2]/K_M, \qquad \alpha \leftarrow \alpha/(K_M\delta), \qquad \beta \leftarrow \beta/(K_M\delta), \qquad K_d \leftarrow K_d/K_M,$$

Here, the quantity to the left of the arrow is the parameter in the non-dimensionalized system. Thus, in the first assignment, the non-dimensionalized time, $t$, is equal to dimensionalized time multiplied by δ.

We can then write the system using the non-dimensionalized quantities:

$$\frac{\delta d K_M[A_{tot}]}{dt} = K_M\delta\alpha + \frac{K_M\delta\beta(K_M[A_2])^n}{K_M^n + (K_M[A_2])^n} - \delta K_M[A_{tot}]$$

$$\frac{\delta d K_M[B_{tot}]}{dt} = K_M\delta\alpha + \frac{K_M\delta\beta(K_M[B_2])^n}{K_M^n + (K_M[B_2])^n} - \delta K_M[B_{tot}]$$

and

$$K_M[A_2] = \frac{2K_M^2[A_{tot}]^2}{\mathrm{K_M}K_d + 4K_M([A_{tot}] + [B_{tot}]) + \sqrt{\mathrm{K_M^2}K_d^2 + 8K_M^2([A_{tot}] + [B_{tot}])K_d}}$$

$$\mathrm{K_M}[B_2] = \frac{2K_M^2[B_{tot}]^2}{\mathrm{K_M}K_d + 4K_M([A_{tot}] + [B_{tot}]) + \sqrt{\mathrm{K_M^2}K_d^2 + 8K_M^2([A_{tot}] + [B_{tot}])K_d}}$$

After canceling δ and $K_M$ from both side of equations, we obtain the non-dimensionalized MultiFate model:

$$\frac{d[A_{tot}]}{dt} = \alpha + \frac{\beta[A_2]^n}{1 + [A_2]^n} - [A_{tot}]$$

$$\frac{d[B_{tot}]}{dt} = \alpha + \frac{\beta[B_2]^n}{1 + [B_2]^n} - [B_{tot}]$$

$$[A_2] = \frac{2[A_{tot}]^2}{K_d + 4([A_{tot}] + [B_{tot}]) + \sqrt{K_d^2 + 8([A_{tot}] + [B_{tot}])K_d}}$$

$$[B_2] = \frac{2[B_{tot}]^2}{K_d + 4([A_{tot}] + [B_{tot}]) + \sqrt{K_d^2 + 8([A_{tot}] + [B_{tot}])K_d}}$$

This non-dimensionalization leaves us with four parameters: rescaled basal protein production rate, α, rescaled maximal protein production rate in the Hill function, β, rescaled dimerization dissociation constant, $K_d$, and Hill coefficient $n$.

Aparts from these four parameters, we used the mathematical model to predict how protein stability control the number of stable fixed points in many parts of this study. In the non-dimensionalized model, the protein degradation rate, δ, does not appear explicitly but enters through the rescaling of α and β by $(\delta K_M)^{-1}$ as shown above. Thus, tuning protein stability is equivalent to multiplying both α and β by a common factor, which we term the "protein stability factor" in this study.

Asymmetric MultiFate-2 model

While the symmetric MultiFate-2 model precisely predicts many experimental results, we observed some asymmetric behaviors in MultiFate-2 lines. For example, MultiFate-2.2 cells in A+B state preferentially migrated towards A-only state when transferred from the High TMP condition to the Low TMP condition (Fig. 3C). This kind of asymmetric behavior could result from several potential differences between integrated TF A cassettes and TF B cassettes: (i) A and B used two different ZF DNA-binding domains (BCRZFR39A and 37ZFR2AR11AR39AR67A). As shown in Fig. S4A, they could have different binding affinity to ZF binding sites, resulting in different $K_M$, and different activated transcriptional rates, resulting in different β; (ii) The integration number of TF A cassettes and TF B cassettes could be different, resulting in different effective values of α and β; (iii) Due to sequence differences in ZF DNA-binding domains, the protein stability could be different for A and B, resulting in different effective values of δ. To analyze such asymmetries, we allow for distinct values of these parameters, indicated by subscripted A or B. While we allow asymmetry in these parameters, we still assume symmetry in others. Specifically, we maintain the same Hill coefficient, $n$, and dissociation constant for dimerization, $K_d$, for both factors because they share the same transcriptional activation domain and the same homodimerization domain. With these assumptions, we can then write down an asymmetric dimensionalized model:

$$\frac{d[A_{tot}]}{dt} = \alpha_A + \frac{\beta_A[A_2]^n}{K_{MA}^n + [A_2]^n} - \delta_A[A_{tot}]$$

$$\frac{d[B_{tot}]}{dt} = \alpha_B + \frac{\beta_B[B_2]^n}{K_{MB}^n + [B_2]^n} - \delta_B[B_{tot}]$$

As above, we then non-dimensionalize the model by rescaling time in units of $\delta_A^{-1}$, and concentrations in units of $K_{MA}$. Then we have (after canceling $K_{MA}$ and $\delta_A^{-1}$ from both sides of the equations):

$$\frac{d[A_{tot}]}{dt} = \alpha_A + \frac{\beta_A[A_2]^n}{1 + [A_2]^n} - [A_{tot}]$$

$$\frac{d[B_{tot}]}{dt} = (\alpha_B/\alpha_A)\alpha_A + \frac{(\beta_B/\beta_A)\beta_A[B_2]^n}{(K_{MB}/K_{MA})^n + [B_2]^n} - \delta_B/\delta_A[B_{tot}]$$

To further simplify these expressions, we define additional parameter ratios, $r = \alpha_B/\alpha_A$, $m = \beta_B/\beta_A$, $\kappa = K_{MB}/K_{MA}$, and $\gamma = \delta_B/\delta_A$. We also let $\alpha = \alpha_A$, and $\beta = \beta_A$. With these definitions, we obtain the ODEs for protein production and degradation in the asymmetric MultiFate model:

$$\frac{d[A_{tot}]}{dt} = \alpha + \frac{\beta[A_2]^n}{1 + [A_2]^n} - [A_{tot}]$$

$$\frac{d[B_{tot}]}{dt} = r\alpha + \frac{m\beta[B_2]^n}{\kappa^n + [B_2]^n} - \gamma[B_{tot}]$$

with the same expressions of the $[A_2]$ and $[B_2]$ in terms of $[A_{tot}]$ and $[B_{tot}]$ shown above.

We now have four parameters that represent different types of asymmetry: $r$ represents the ratio of basal protein production rates, $m$ represents the ratio of maximal protein production rates by self-activation, κ represents the ratio of homodimer concentrations for half-maximal activation, and γ represents the ratio of protein degradation rates. The symmetric MultiFate-2 model is then a special case where $r = m = \kappa = \gamma = 1$.

MultiFate model expanded to N transcription factors

The minimal MultiFate-2 model can be expanded in a straightforward way to include more transcription factors. To start, we consider the distribution of transcription factors $X_1$, $X_2$, $X_3$, … $X_N$ among different dimerization states. For each transcription factor, the total concentration can be expressed as,

$$[X_{tot,i}] = [X_i] + \sum_{j \neq i} [X_i][X_j] + 2[X_{2,i}]$$

$$for\ i\ = 1, 2, 3, \ldots\ N$$

Here $[X_i]$ denotes the concentration of transcription factor $X_i$ monomers, $[X_{2,i}]$ denote the concentration of homodimers, and $[X_i][X_j]$ denote the concentration of heterodimers formed by $X_i$ and $X_j$ ($i \neq j$). As mentioned above, we assume that homo- and hetero-dimerization occur with equal dissociation constants, $K_d$, reflecting the use of the same dimerization domain for both proteins. Because dimerization dynamics occur on a faster timescale than protein production and degradation, the protein dimerization states approximately follow their equilibrium values:

$$[X_i]^2 = K_d[X_{2,i}]$$

$$2[X_i][X_j] = K_d[X_i][X_j]$$

$$for\ i = 1, 2, 3, \ldots\ N\ and\ i \neq j$$

Solving these equations produces a simple expression for the concentrations of the activating homodimers in terms of the total concentrations of all transcription factor species:

$$[X_{2,i}] = \frac{2[X_{tot,i}]^2}{K_d + 4\sum[X_{tot,i}] + \sqrt{K_d^2 + 8K_d \sum[X_{tot,i}]}}$$

$$for\ i\ = \ 1, 2, 3, \ldots\ N$$

With these expressions, we can describe protein production and degradation dynamics using ODEs for $[X_{tot,i}]$ in a similar way as shown in Box 1. After non-dimensionalization and adding asymmetric parameters, we obtain ODEs for protein production and degradation in the expanded MultiFate model:

$$\frac{d[X_{tot,i}]}{dt} = r_i\alpha + \frac{m_i\beta[X_{2,i}]^n}{\kappa_i^n + [X_{2,i}]^n} - \gamma_i[X_{tot,i}]$$

$$for\ i\ = \ 1, 2, 3, \ldots\ N$$

where $\alpha$ represents the basal protein production, $\beta$ represents the maximal protein production rate in the Hill function, $n$ represents Hill coefficient and $r_i$, $m_i$, $\kappa_i$, and $\gamma_i$ represents the asymmetric parameters for transcription factor $X_i$.

<u>MultiFate-2 model with mRNA and protein dimerization dynamics</u>

The treatment above lumps the processes of mRNA transcription and protein translation together into a single gene expression step and uses a steady-state approximation for dimerization dynamics. This model works accurately to predict the number and locations of stable fixed points. However, to capture the dynamics of cells during bifurcation and state-switching events, an expanded model that includes both mRNA and protein dimerization dynamics is required.

To incorporate mRNA dynamics into the model, we assume TF A and TF B mRNAs, denoted $[a]$ and $[b]$, are produced at a total rate equal to their basal transcription rate $k_1$ plus a homodimerdependent transcriptional activation rate, which follows a Hill function of corresponding homodimer concentration, $[A_2]$ and $[B_2]$, with maximal rate $k_2$, Hill coefficient $n$, and half-maximal activation at a homodimer concentration of $K_M$. Each mRNA species is removed at a total rate $\delta_{mRNA}$. For generality, we also allow asymmetry parameters, with $r$ representing the ratio between A and B basal transcription rates ($r = k_{1,a}/k_{1,b}$), $m$ representing the ratio between A and B maximal rates ($m = k_{2,a}/k_{2,b}$), and κ representing the ratio between A and B half-maximal concentrations ($\kappa = K_{M,a}/K_{M,b}$). We can then write:

$$\frac{d[a]}{dt} = k_1 + \frac{k_2[A_2]^n}{K_M^n + [A_2]^n} - \delta_{mRNA}[a]$$

$$\frac{d[b]}{dt} = rk_1 + \frac{mk_2[B_2]^n}{(\kappa K_M)^n + [B_2]^n} - \delta_{mRNA}[b]$$

Where we just let $k_1 = k_{1,a}$ and $k_2 = k_{2,a}$ for notational simplicity.

Next, we describe the dynamics of TF A and TF B proteins in different dimerization forms with ODEs. For monomers of TF A and TF B, denoted $[A]$ and $[B]$, each equation consists of terms describing translation, protein removal, monomer association, dimer dissociation or conversion due to degradation of one of the constituent monomers. Here $k_p$ is the translation rate, $\delta$ is the protein removal rate, $k_{on}$ is the monomer association rate and $k_{off}$ is the dimer dissociation rate. $[AB]$ denotes the concentration of AB heterodimers.

The asymmetry parameter represents the ratio of protein A and B removal rates, $\gamma = \delta_A/\delta_B$, where we just let $\delta = \delta_A$ for notational simplicity:

$$\frac{d[A]}{dt} = k_p[a] - \delta[A] - 2k_{on}([A]^2 + [A][B]) + k_{off}(2[A_2] + [AB]) + \gamma\delta[AB]$$

$$\frac{d[B]}{dt} = k_p[b] - \gamma\delta[B] - 2k_{on}([B]^2 + [B][A]) + k_{off}(2[B_2] + [AB]) + \gamma\delta[AB]$$

For dimers, each equation consists of terms for removal, association, and dissociation:

$$\frac{d[A_2]}{dt} = -\delta[A_2] + 2k_{on}[A]^2 - k_{off}[A_2]$$

$$\frac{d[B_2]}{dt} = -\gamma\delta[B_2] + 2k_{on}[B]^2 - k_{off}[B_2]$$

$$\frac{d[AB]}{dt} = -\delta[AB] - \gamma\delta[AB] + 2k_{on}[A][B] - k_{off}[AB]$$

Stochastic modeling of MultiFate circuits

To simulate the dynamics of the MultiFate-2 system during state-switching (Fig. 3F and Movie S1) and bifurcation (Fig. 3E and Movie S3) events, we constructed a stochastic model based on the same reactions represented by the ODEs in the above MultiFate-2 model with mRNA and protein dimerization dynamics. Molecular reactions and their propensities for Gillespie simulation (*88*) are listed in Table S3. All terms have a concentration unit of molecule number per cell (either mRNA or protein), and a time unit of hours. We performed Gillespie simulations using the biocircuits Python package (https://pypi.org/project/biocircuits/) with physiologically reasonable parameters (see below).

Physiologically reasonable parameter regimes

From existing literature and experimental measurements performed in this study, we estimated the physiologically reasonable regime for each dimensionalized parameter ($K_M$, δ, α, β, $K_d$, $n$, $k_1$, $k_2$, $\delta_{mRNA}$, $k_p$, $k_{on}$, $k_{off}$). Estimated values for these parameters are summarized in Table S4.

Since some measurement data are in the unit of concentration, while others are in the unit of molecules per cell, we first estimated the number of molecules equivalent to 1 nM in a CHO-K1 cell. The diameter of a CHO cell is ~14 µm (*90*), from which we can calculate the cell volume to be around $1.4\times10^{-12}$ L (assuming it to be a sphere). With this rough approximation, **1 nM = 1 nM $\times$ $1.4\times10^{-12}$ L $\times$ $6\times10^{23}$ molecules/mol ≈ 800 molecules per CHO cell.** Below, we used this value to convert between molecules per cell and molarity.

The concentration for half-maximal activation, $K_M$, of our ZF activator homodimer, is used to rescale all concentrations in the non-dimensionalized MultiFate model (see above). We could not find a direct *in vivo* measurement of this value in the literature. However, the $K_M$ of a monomeric ZF activator with a 3-finger Zif268 ZF domain was estimated to be ~600 nM (assuming the volume of yeast cells to be 40 µm$^3$) (*91*, *92*). Another *in vitro* study showed that by linking two Zif268 ZF domain with a linker, the resulting 6-finger ZF domain could bind to a 18bp DNA target site almost 70-fold stronger than a single Zif268 domain binding to a 9bp target site (*93*). Although *in vivo* and the *in vitro* $K_M$ can differ by several orders of magnitudes due to the interactions of proteins with genomic background, the fold changes in $K_M$ are in reasonable agreement between the *in vivo* and the *in vitro* results (*94*). Therefore, we estimated that an activator with a 6-finger ZF domain has a KM of 600 nM / 70 ≈ 8 nM. Our ZF transcription factor homodimer should bind to 18bp target DNA site in a similar fashion with how 6-finger ZF domain does, thus we estimated our $K_M$ to be comparable to, or (due to a more complicated genomic background in mammalian cells), slightly larger than the range of 8 – 20 nM. Based on this reasoning, **we used a $K_M$ = 10 nM = 8000 molecules/cell in the model.**

Next, we estimated parameters related to protein production and removal dynamics. We used a protein removal rate δ = 0.1 hr$^{-1}$ for stable proteins in our system, based on an *in vivo* measurement of proteome half-life dynamics in living human cells (*95*). Our engineered ZF transcription factors have DHFR domains at their C-terminus, whose protein removal rate is controlled by TMP concentration. The dynamic range of this regulation is at least 20 fold (*61*, *96*). Therefore, we estimated the range of δ to be between 0.1 hr$^{-1}$ and 2 hr$^{-1}$, under saturating TMP condition and no TMP condition, respectively. **In the model, we used δ = 0.1 hr$^{-1}$ for the "High TMP" condition, and δ = 0.2 hr$^{-1}$ for the "Low TMP" condition.**

We next estimated $k_2$, $\delta_{mRNA}$ and $k_p$, which together are critical for establishing the levels and dynamics of mRNA and protein. In mammalian cells, average transcription rates were estimated to be ~2 mRNA/(gene•hr) (*97*). Based on previous stable cell line construction using PiggyBac in our lab (*98*), we estimated that a total of 50–100 gene cassettes were integrated during the construction of MultiFate-2, with 25–50 copies integrated each, for TF A and TF B. Maximal transcription rate $k_2$ was then estimated to be 50–100 mRNA/(cell•hr), and **we used an intermediate value $k_2$ = 80 mRNA/(cell•hr).** For typical mRNA half-life, different studies have provided diverse values, ranging from 50 minutes to 9 hours (*97*, 99, *100*). This corresponds to a mRNA removal rate mRNA ranging from ln(2) / (9 hr) – ln(2) / (50 min) ≈ 0.077 – 0.83 $hr^{-1}$. Since these studies did not take mRNA dilution from cell division into consideration, **we used a value of $\delta_{mRNA}$ = 0.7 $hr^{-1}$,** closer to the upper bound of the estimated range, in our stochastic model. For protein translation rate, **we used a value of $k_p$ = 140 proteins/(mRNA•hr)** based on one of the above studies (*97*).

The value of β can be estimated from the above parameters. Since the mRNA removal rate is much higher than the protein removal rate, we assumed that mRNA dynamics is approximately at steady state on the timescale of $\delta^{-1}$. This assumption enabled us to estimate the **maximal protein production rate in the Hill function as $\beta = k_2 \times k_p/\delta_{mRNA}$ = 16000 proteins/(cell•hr) = 20 nM/hr.** In the experiment, we observed that there was a fluorescence expression difference of around 25-50 fold between ON and OFF states (Fig. 3B). (The estimate of the OFF level is not limited by autofluorescence). Since expression in the OFF state comes from basal transcription, we estimated the basal transcription rate $k_1$ to be 25–50 fold smaller than $k_2$, giving a range of 1.152–4.608 mRNA/hr. From this, **we used an intermediate value of $k_1 =$ 3.2 mRNA/(cell•hr). Similarly, we can estimate the basal protein production rate $\alpha = k_1 \times k_p/\delta_{mRNA}$ = 640 proteins/(cell•hr) = 0.8 nM/hr.**

The final parameter related to protein production dynamics is the Hill coefficient $n$. Different studies obtained very different measurements of transcriptional Hill coefficients (*91*, *101–105*), ranging from 1–3.6. **Here we used a modest Hill coefficient of $n$ = 1.5.**

Finally, we estimated parameters related to protein dimerization. The apparent dissociation constant $K_d$ of FKBP homodimerization domain should depend on AP1903 concentrations. However, we could not find a direct measurement of this dependency. Therefore, we compared FKBP with another homodimerization domain GCN4 that we used in this study (Fig. 2A), which was shown to have a $K_d$ of 10–20 nM (*106*). ZF transcription factors with FKBP (Fig. 2B) more strongly activated the reporter in 100 nM AP1903 media than ZF transcription factor with GCN4 did (Fig. S4B, BCRZF). Based on this observation, we reasoned that in media containing 100 nM AP1903, the apparent dissociation constant $K_d \leq$ 10 nM. **We therefore used an estimate of $K_d$ = 10 nM = 8000 molecules/cell in the model. For monomer association rate $k_{on}$, we used an intermediate value in the range of diffusion-limited association rates of $k_{on}$ = 4x10$^5$/(M•s) = 1.8x10$^{-3}$ cell/(protein•hr)** (*107*)**.** These two values together produce a **dimer dissociation rate $k_{off} = K_d \times k_{on}$ = 14.4 $hr^{-1}$.**

Source of ultrasensitivity in MultiFate circuit

To generate multistability, a circuit should have both positive feedback and some levels of ultrasensitivity (i.e. effective Hill exponent greater than 1) (*108–110*). Although transcriptional activation can exhibit some ultrasensitivity in mammalian cells (represented by the Hill coefficient of $n = 1.5$ above) (*101*, *111*), parameter screening revealed that MultiFate generates multistability even when $n = 1$ (no ultrasensitivity from transcriptional activation). This provokes the question of where the additional ultrasensitivity comes from in the MultiFate circuit. Two features of the MultiFate circuit could provide this additional ultrasensitivity. First, transcription factors homodimerize to self-activate, and homodimerization has been shown to introduce ultrasensitivity (*112*, *113*). Indeed, dimerization results in a $[A_{tot}]^2$ (or $[B_{tot}]^2$) term in the numerator of the expression for $[A_2]$ (or $[B_2]$) in Box 1, which we write again here for convenience:

$$[A_2] = \frac{2[A_{tot}]^2}{K_d + 4([A_{tot}] + [B_{tot}]) + \sqrt{K_d^2 + 8([A_{tot}] + [B_{tot}])K_d}}$$

$$[B_2] = \frac{2[B_{tot}]^2}{K_d + 4([A_{tot}] + [B_{tot}]) + \sqrt{K_d^2 + 8([A_{tot}] + [B_{tot}])K_d}}$$

Its contribution to ultrasensitivity depends on the dimerization dissociation constant $K_d$. When dimerization is strong (small $K_d$), the $4([A\_\{tot\}] + [B\_\{tot\}]$ term dominates in the denominator, which cancels with the quadratic term $[A_{tot}]^2$ or $[B_{tot}]^2$ in the numerator. This makes the expression more linear, thus reducing the ultrasensitivity by homodimerization. Conversely, when dimerization is weak (large $K_d$), the $K_d^2$ term dominates in the denominator, which makes the expression more quadratic and increases the ultrasensitivity by homodimerization.

A second source of ultrasensitivity comes from mutual inhibition through heterodimerization, a prevalent feature in biology also known as molecular titration, which has been shown to introduce ultrasensitivity (*109*, *112*). Here, opposite to the case with homodimerization, strong dimerization (small $K_d$) increases the ultrasensitivity introduced through molecular titration (*112*). Together, additional ultrasensitivity comes mainly from homodimerization when $K_d$ is large, and mainly from molecular titration when $K_d$ is small. This explains why the MultiFate circuit generates multistability in a wide $K_d$ range (Fig. S1 and Fig. S2).

Modulating leaky expression by modifying synthetic promoter sequences

When characterizing different ZF transcription factors, we found that promoters containing the 9bp binding site GACGCTGCT for 42ZF (*52*) have higher basal expression. Basal protein production of a cassette then can be modulated by introducing different numbers of GACGCTGCT motifs at promoter regions.

### Fig. S1

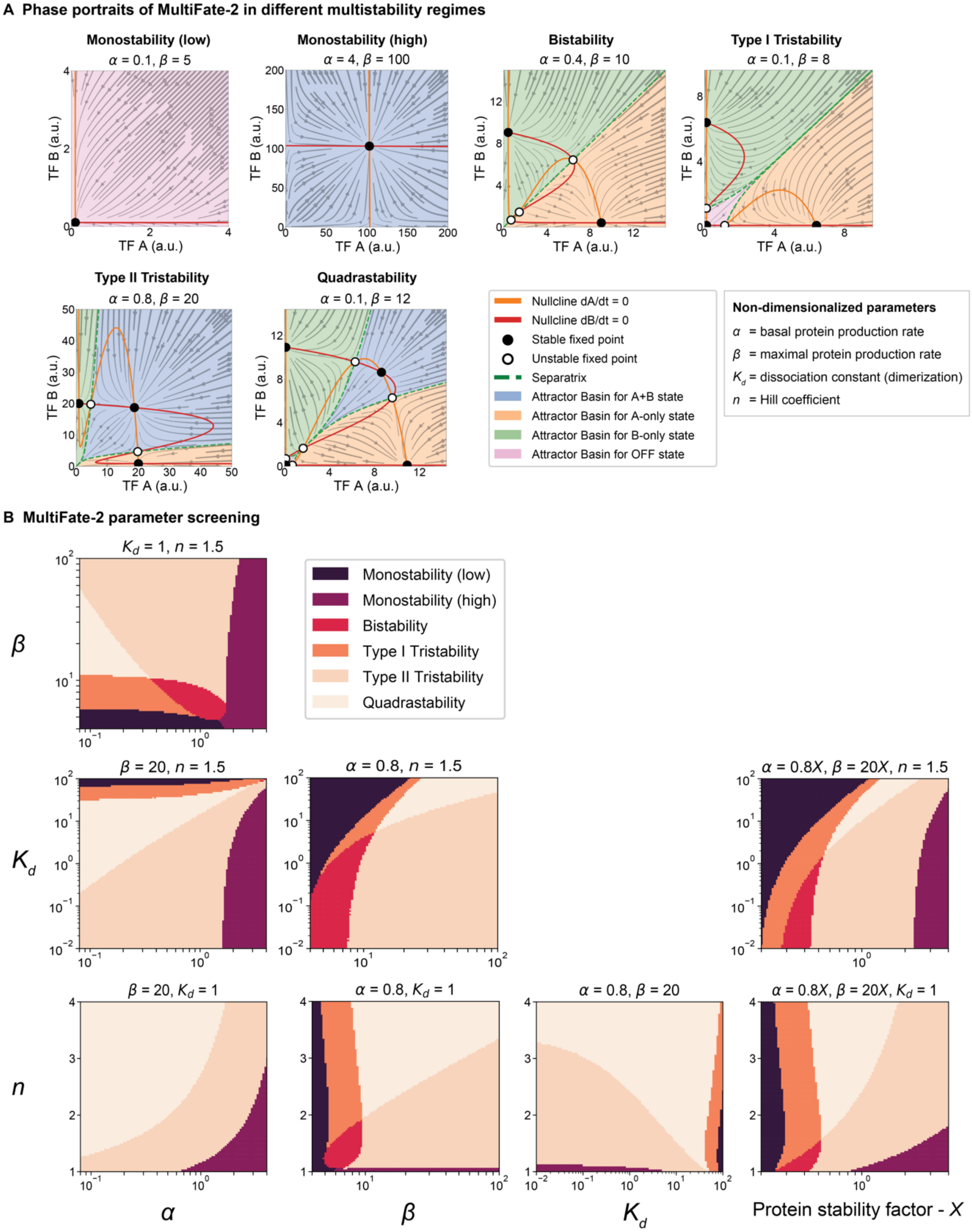
A Phase portraits of MultiFate-2 in different multistability regimes
Monostability (low)
α = 0.1, β = 5
Monostability (high)
α = 4, β = 100
Bistability
α = 0.4, β = 10
Type I Tristability
α = 0.1, β = 8
Type II Tristability
α = 0.8, β = 20
Quadrastability
α = 0.1, β = 12
TF A (a.u.)
TF B (a.u.)
Nullcline dA/dt = 0
Nullcline dB/dt = 0
Stable fixed point
Unstable fixed point
Separatrix
Attractor Basin for A+B state
Attractor Basin for A-only state
Attractor Basin for B-only state
Attractor Basin for OFF state
Non-dimensionalized parameters
α = basal protein production rate
β = maximal protein production rate
$K_d$ = dissociation constant (dimerization)
n = Hill coefficient
B MultiFate-2 parameter screening
$K_d$ = 1, n = 1.5
Monostability (low)
Monostability (high)
Bistability
Type I Tristability
Type II Tristability
Quadrastability
β = 20, n = 1.5
α = 0.8, n = 1.5
α = 0.8X, β = 20X, n = 1.5
β = 20, $K_d$ = 1
α = 0.8, $K_d$ = 1
α = 0.8, β = 20
α = 0.8X, β = 20X, $K_d$ = 1
α
β
$K_d$
n
Protein stability factor - X

**Fig. S1. The MultiFate-2 model generates diverse types of multistability.**

(A) In different symmetric parameter regimes (in which parameters for the two transcription factors are identical), MultiFate-2 can generate two types of monostability, bistability, two types of tristability, and quadrastability. For each regime, non-dimensionalized parameters α and β are provided above the plot, and $K_d = 1$ and $n = 1.5$. (B) Parameter screen reveals how each of the non-dimensionalized parameters individually affects the global structure of the system. Each row and column in this grid of plots represents a titration of one parameter value, indicated at left and bottom. Within each plot, different colors represent different stability regimes, as in (A), determined by numerically solving for steady state values and their linear stability at each point in each parameter space (Supplementary Methods). In the non-dimensionalized model, changing protein stability is equivalent to multiplying α and β with the same factor, and is shown in the fourth column ("protein stability factor - X"), with higher values representing greater protein stability. Higher leaky transcription (high α) allows transcription factors to self-activate, destabilizing the OFF state (α column of plots). Very high α values push the system towards monostability where only the state in which both transcription factors are highly expressed is stable. Stronger self-activation (higher values of β) is more likely to produce type II tristability and quadrastability (β column of plots). Strong dimerization (low $K_d$) is essential for type II tristability ($K_d$ row of plots). A broad range of Hill coefficients $n \geq 1$ are compatible with different types of multistability ($n$ row of plots). While higher values of $n$ reduce sensitivity to other parameters and allow the system generate type II tristability even with higher values of α, they also stabilize the OFF state to favor quadrastability.

### Fig. S2

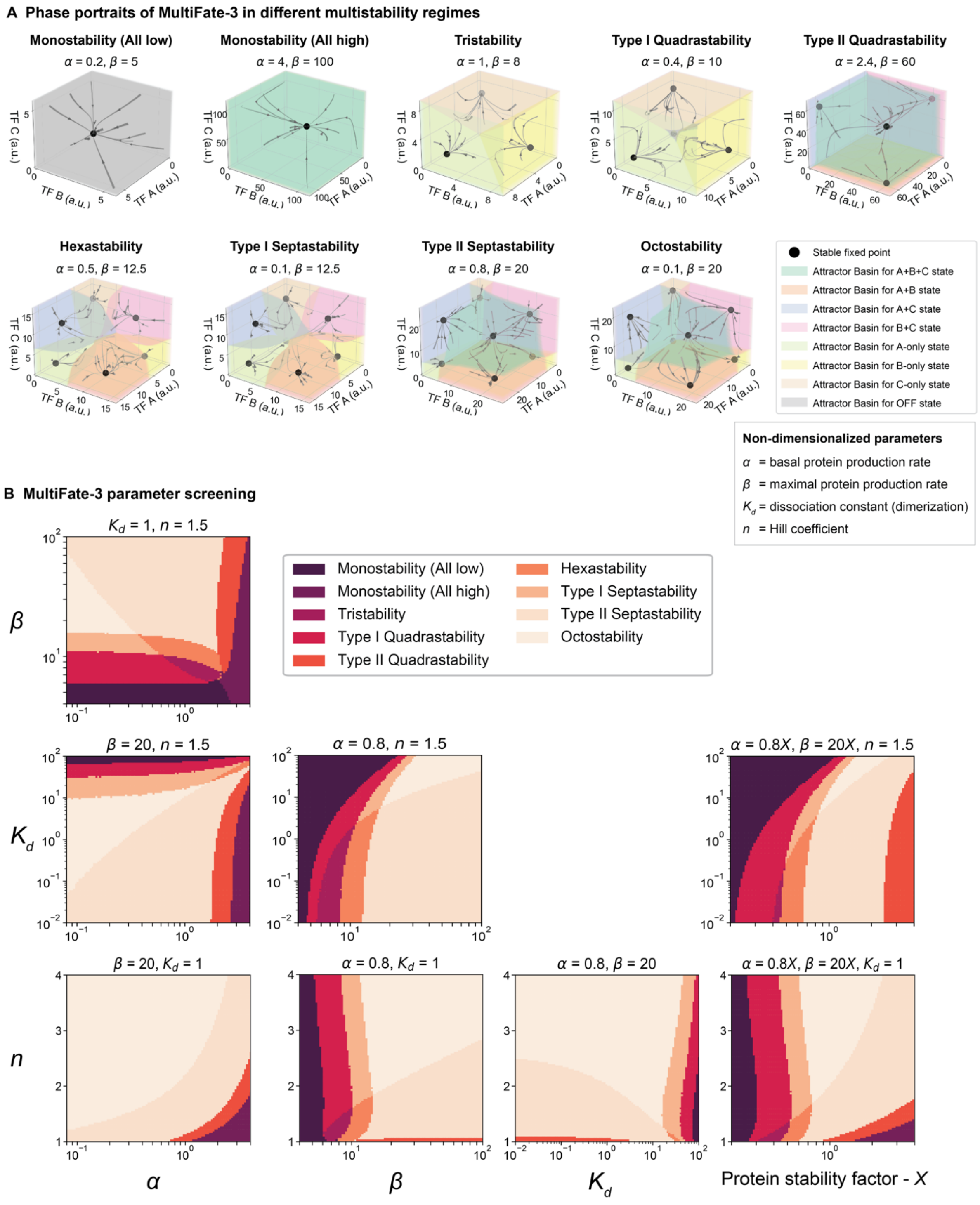
A Phase portraits of MultiFate-3 in different multistability regimes
Monostability (All low)
α = 0.2, β = 5
Monostability (All high)
α = 4, β = 100
Tristability
α = 1, β = 8
Type I Quadrastability
α = 0.4, β = 10
Type II Quadrastability
α = 2.4, β = 60
Hexastability
α = 0.5, β = 12.5
Type I Septastability
α = 0.1, β = 12.5
Type II Septastability
α = 0.8, β = 20
Octostability
α = 0.1, β = 20
TF A (a.u.)
TF B (a.u.)
TF C (a.u.)
Stable fixed point
Attractor Basin for A+B+C state
Attractor Basin for A+B state
Attractor Basin for A+C state
Attractor Basin for B+C state
Attractor Basin for A-only state
Attractor Basin for B-only state
Attractor Basin for C-only state
Attractor Basin for OFF state
Non-dimensionalized parameters
α = basal protein production rate
β = maximal protein production rate
Kd = dissociation constant (dimerization)
n = Hill coefficient
B MultiFate-3 parameter screening
Kd = 1, n = 1.5
β = 20, n = 1.5
α = 0.8, n = 1.5
α = 0.8X, β = 20X, n = 1.5
β = 20, Kd = 1
α = 0.8, Kd = 1
α = 0.8, β = 20
α = 0.8X, β = 20X, Kd = 1
Monostability (All low)
Monostability (All high)
Tristability
Type I Quadrastability
Type II Quadrastability
Hexastability
Type I Septastability
Type II Septastability
Octostability
α
β
Kd
n
Protein stability factor - X

**Fig. S2. The MultiFate-3 model generates more diverse types of multistability compared with MultiFate-2.**

(A) In different parameter regimes, MultiFate-3 can generate a diverse repertoire of different types of stability. These include two types of monostability, tristability, two types of quadrastability, hexastability, two types of septastability, and octastability. For each regime, non-dimensionalized parameter values are provided above the plot (Box 1). Movie S4 provides a rotating 3-dimensional view of all of these phase diagrams. For each regime, non-dimensionalized parameters α and β are provided above the plot, and $K_d = 1$ and $n = 1.5$. (B) Parameter screen reveals how each of the non-dimensionalized parameters individually affects the global structure of the system. As in Fig. S1, changing protein stability in the non-dimensionalized model is equivalent to multiplying α and β with the same factor, denoted "protein stability factor - X," with higher values representing greater protein stability. Each plot in the grid shows a titration of two parameter values (left and bottom). Higher leaky transcription (high α) allows transcription factors to self-activate, destabilizing the OFF state. Very high α pushes the system towards monostability where only the state which all transcription factors are highly expressed is stable (α column of plots). Stronger self-activation (higher values of β) generally favors higher levels of multistability, such as septastability and octostability (β column of plots). Strong dimerization (low $K_d$) is essential for type II septastability ($K_d$ column of plots). A broad range of Hill coefficients $n \geq 1$ are compatible with different types of multistability. Higher values of $n$ reduce sensitivity to other parameters, and allow the system to generate type II septastability even at higher values of α. However, they also stabilize the OFF state to favor octostability ($n$ column of plots). Note that the MultiFate-3 parameter screen graph structure resembles that of MultiFate-2 (Fig. S1B). Two types of monostability for both MultiFate-2 and MultiFate-3 appear at similar positions. Octostability in MultiFate-3 occurs at a similar position as quadrastability in MultiFate-2. Similarly, type II septastability in MultiFate-3 is at a similar position as type II tristability in MultiFate-2. As in Fig. S1B, each plot is calculated by numerical solution of the MultiFate-3 model for steady state values and their linear stability (Supplementary Materials).

### Fig. S3

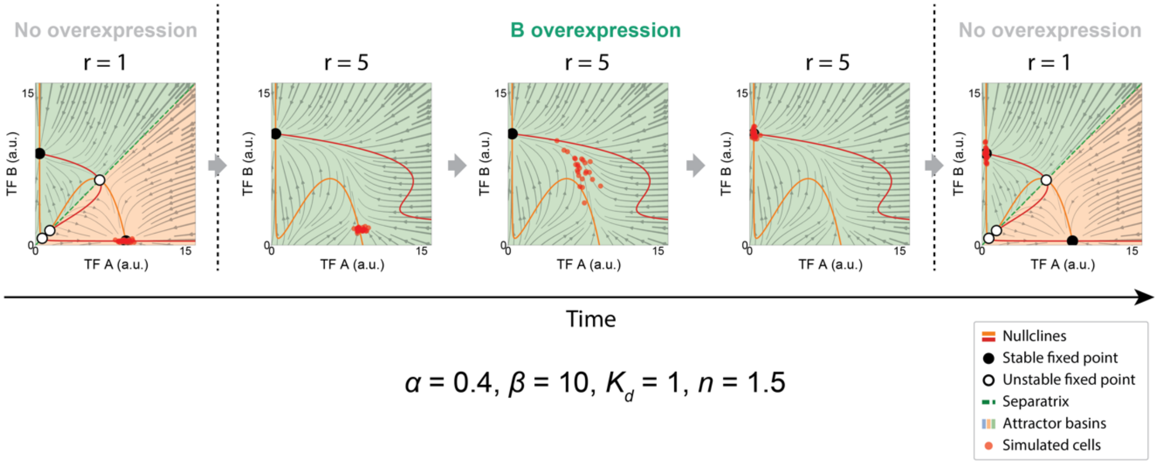


$\alpha = 0.4$, $\beta = 10$, $K_d = 1$, $n = 1.5$

**Fig. S3. Modeling state-switching dynamics.**

Here, we show the effect of transiently expressing additional B on switching of cells from the A-only state to the B-only state in the bistable regime (Fig. S1). Ectopic B expression is represented by the parameter $r$ (Supplementary Materials), defined as the basal protein production rate of B divided by the basal protein production rate of A, which quantifies the transient asymmetry in the model. Initially (left plot), cells (red dots) are in the A-only state. Increasing $r$ destabilizes the A-only state but not the B-only state (central three plots), allowing cells to migrate to the B-only state. Terminating ectopic B expression restores $r = 1$. However, at this point, cells are already stabilized in the B-only attractor. These plots are shown as an animation in Movie S1.

### Fig. S4

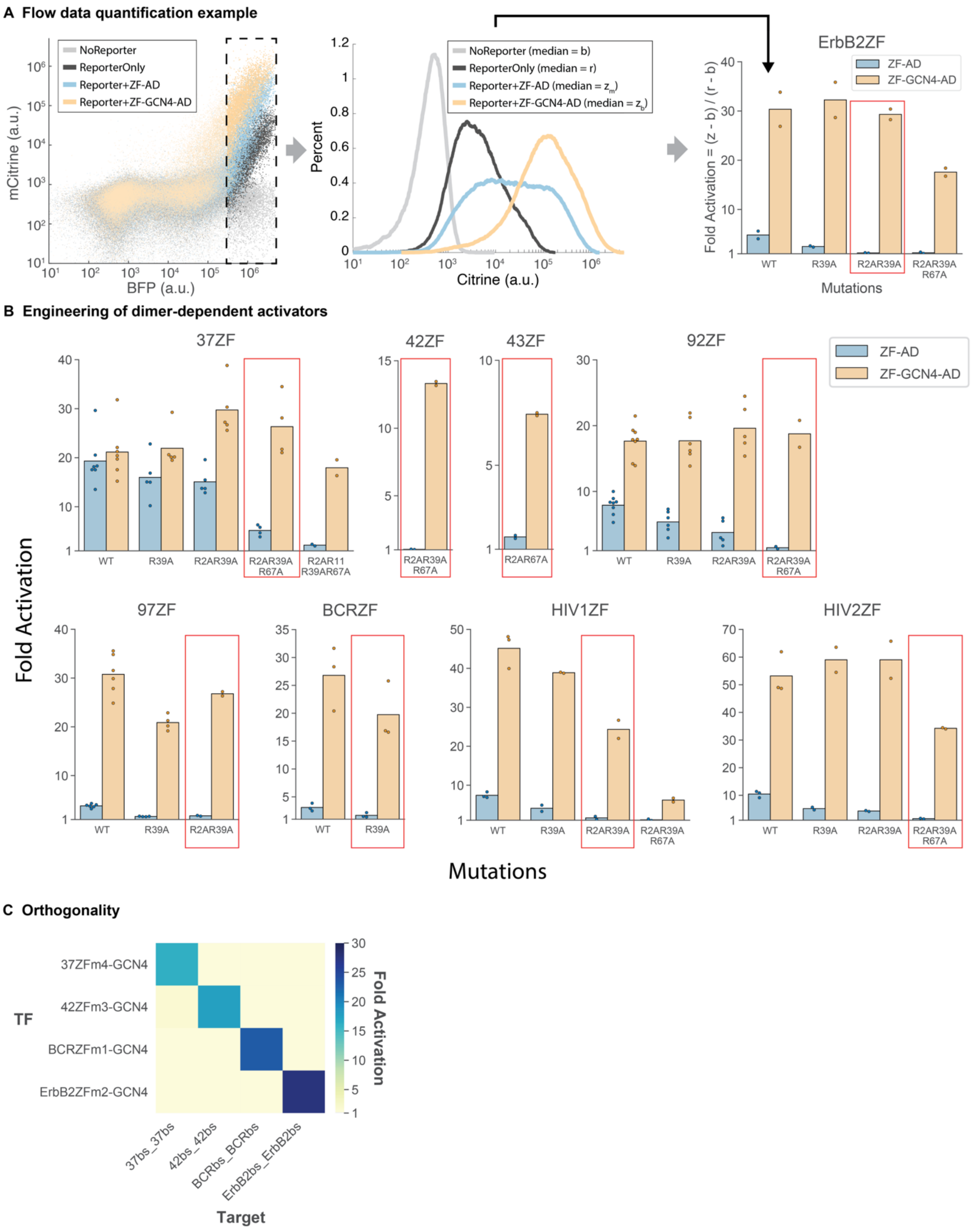

A Flow data quantification example
NoReporter
ReporterOnly
Reporter+ZF-AD
Reporter+ZF-GCN4-AD
mCitrine (a.u.)
BFP (a.u.)
Percent
NoReporter (median = b)
ReporterOnly (median = r)
Reporter+ZF-AD (median = $z_m$)
Reporter+ZF-GCN4-AD (median = $z_b$)
Citrine (a.u.)
ErbB2ZF
ZF-AD
ZF-GCN4-AD
Fold Activation = (z - b) / (r - b)
WT
R39A
R2AR39A
R2AR39A R67A
Mutations
B Engineering of dimer-dependent activators
37ZF
42ZF
43ZF
92ZF
ZF-AD
ZF-GCN4-AD
Fold Activation
WT
R39A
R2AR39A
R2AR39A R67A
R2AR11 R39AR67A
R2AR67A
97ZF
BCRZF
HIV1ZF
HIV2ZF
Mutations
C Orthogonality
TF
37ZFm4-GCN4
42ZFm3-GCN4
BCRZFm1-GCN4
ErbB2ZFm2-GCN4
37bs_37bs
42bs_42bs
BCRbs_BCRbs
ErbB2bs_ErbB2bs
Target
Fold Activation

**Fig. S4. Engineering dimer-dependent transcriptional regulation.**

(A) To characterize the activation strength of different zinc finger transcription factor variants, we co-transfected each of them with a reporter construct (Citrine) and a co-transfection marker (mTagBFP2). Highly transfected cells were gated with co-transfected BFP > $3x10^5$ (dashed box) to extract their individual histograms (middle panel). From each histogram, median Citrine fluorescence intensities of gated cells was used to calculate fold activation (right panel, Supplementary Materials). Black arrow indicates the original zinc finger sequence from the middle panel (WT). The R2AR39A variant (red box) was selected for its high ZF-GCN4-AD activation and minimal ZF-AD activation (AD denotes the VP48 activation domain). (B) Zinc finger mutation variants with minimal activation by ZF-AD and strong activation by ZF-GCN4-AD (red boxes) were selected for use in MultiFate circuits. Each dot represents one replicate, and each bar indicates the mean of all replicates. 37ZF, 42ZF, 43ZF, 92ZF, 97ZF were taken from (*52*). These zinc fingers were previously (*52*) denoted, respectively, 37-12 array, 42-10 array, 43-8 array, 92-1 array, and 97-4 array. BCRZF, HIV1ZF, HIV2ZF, and ErbB2ZF are from (*56*), while BCR denotes the BCR_ABL domain. (C) Four selected zinc finger transcription factors used later in the paper exhibit orthogonal trans-activation. Each row represents one transcription factor and each represents a target reporter construct. Labels for the rows are abbreviated for simplicity. Full transcription factor descriptions are, from top to bottom, 37ZFR2AR11AR39AR67A-GCN4-VP48; 42ZFR2AR39AR67A-GCN4-VP48; BCRZFR39A-GCN4-VP48; and ErbB2ZFR2AR39A-GCN4-VP48. Each target (column) is the same Citrine fluorescent reporter used in panel (A), with 2 repeats of 18bp tandem binding site pairs, denoted "ZFbs_ZFbs" for each type, at the promoter. Each square in the matrix is the mean of two replicates.

Fig. S5

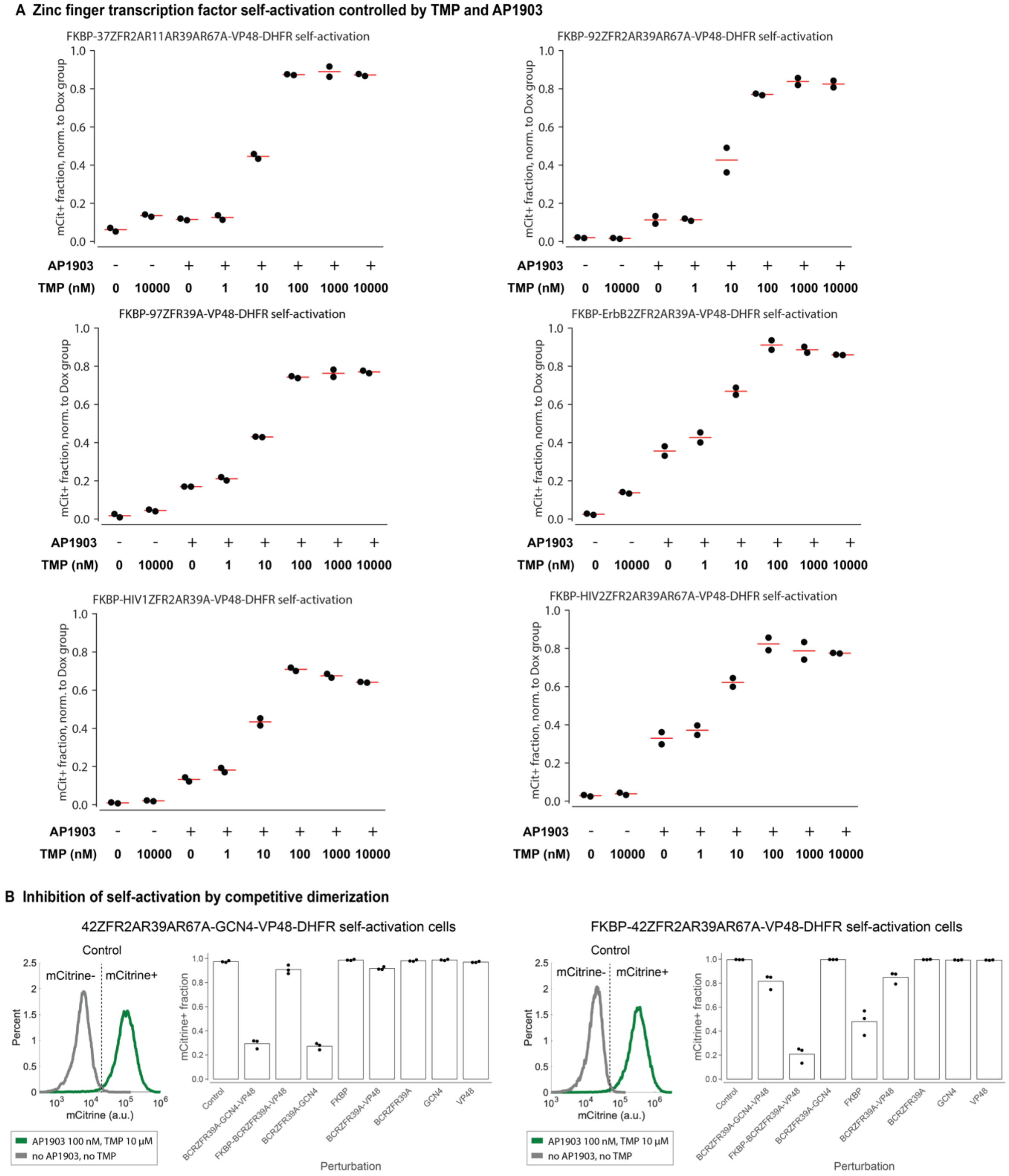

A Zinc finger transcription factor self-activation controlled by TMP and AP1903
FKBP-37ZFR2AR11AR39AR67A-VP48-DHFR self-activation
FKBP-92ZFR2AR39AR67A-VP48-DHFR self-activation
FKBP-97ZFR39A-VP48-DHFR self-activation
FKBP-ErbB2ZFR2AR39A-VP48-DHFR self-activation
FKBP-HIV1ZFR2AR39A-VP48-DHFR self-activation
FKBP-HIV2ZFR2AR39AR67A-VP48-DHFR self-activation
mCit+ fraction, norm. to Dox group
AP1903
TMP (nM)
0 10000 0 1 10 100 1000 10000
B Inhibition of self-activation by competitive dimerization
42ZFR2AR39AR67A-GCN4-VP48-DHFR self-activation cells
FKBP-42ZFR2AR39AR67A-VP48-DHFR self-activation cells
Control
mCitrine-
mCitrine+
Percent
mCitrine (a.u.)
AP1903 100 nM, TMP 10 μM
no AP1903, no TMP
mCitrine+ fraction
Control
BCRZFR39A-GCN4-VP48
FKBP-BCRZFR39A-VP48
BCRZFR39A-GCN4
FKBP
BCRZFR39A-VP48
BCRZFR39A
GCN4
VP48
Perturbation

**Fig. S5. Engineered dimer-dependent transcription factors enable transcriptional positive autoregulation and mutual inhibition through competitive dimerization.**

(A) Positive autoregulation can be controlled by TMP and AP1903. Here, normalized mCitrine+ fractions are used to quantify self-activation strength, as in Fig. 2C (see Supplementary Materials for quantification methods) for 6 additional ZFs. (B) Positive autoregulation was inhibited by competing proteins with matching dimerization domains. Two monoclonal stable lines (plot subtitles) could spontaneously self-activate in media containing AP1903 and TMP (histograms). Potentially competing proteins were expressed from plasmids and stably integrated into each monoclonal line (Supplementary Materials). Note that in 42ZFR2AR39AR67A-GCN4-VP48-DFHR self-activation cells, the GCN4 domain by itself did not have inhibitory effects, but could efficiently inhibit self-activation when fused with BCRZFR39A. In FKBP-42ZFR2AR39AR67A-VP48-DHFR self-activation cells, the FKBP domain by itself can partially inhibit self-activation. The mCitrine threshold is $2x10^4$ for 42ZFR2AR39AR67A-GCN4-VP48-DHFR self-activating cells (left), and $5x10^4$ for FKBP-42ZFR2AR39AR67A-VP48-DHFR cells (right). In all panels, each dot represents one replicate, and each red line or bar indicates the mean of replicates.

### Fig. S6

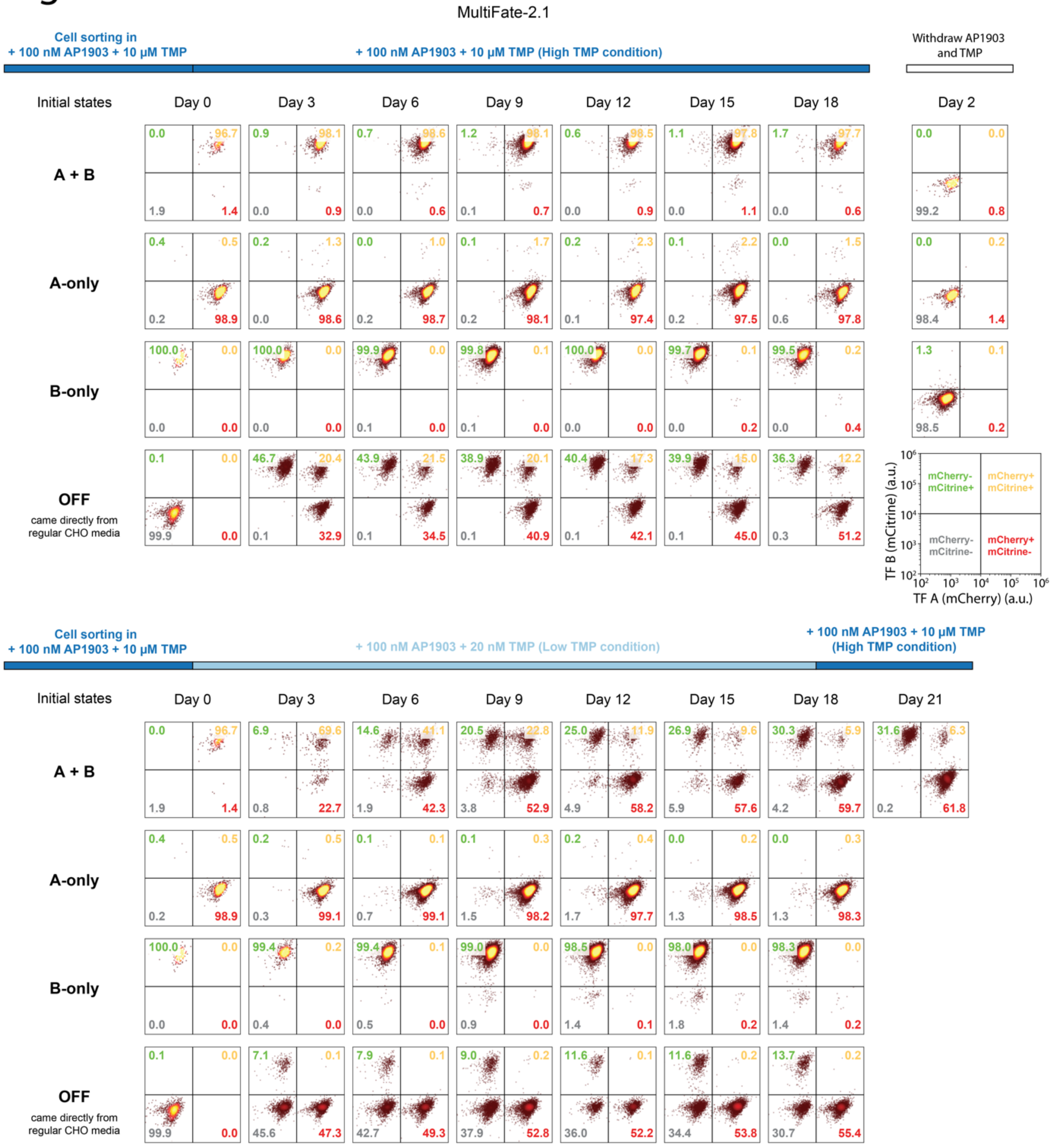
MultiFate-2.1
Cell sorting in
+ 100 nM AP1903 + 10 μM TMP
+ 100 nM AP1903 + 10 μM TMP (High TMP condition)
Withdraw AP1903 and TMP
Initial states
Day 0
Day 3
Day 6
Day 9
Day 12
Day 15
Day 18
Day 2
A + B
A-only
B-only
OFF
came directly from regular CHO media
mCherry- mCitrine+
mCherry+ mCitrine+
mCherry- mCitrine-
mCherry+ mCitrine-
TF B (mCitrine) (a.u.)
TF A (mCherry) (a.u.)
Cell sorting in
+ 100 nM AP1903 + 10 μM TMP
+ 100 nM AP1903 + 20 nM TMP (Low TMP condition)
+ 100 nM AP1903 + 10 μM TMP (High TMP condition)
Day 21

**Fig. S6. Raw flow cytometry analysis of the MultiFate-2.1 line.**

Each plot represents one of three replicates at the indicated time point (cf. Fig. 3C, E). Initial A-only, B-only and A+B cells (rows) were sorted under the media conditions indicated on the top, and initial OFF cells came directly from cells in regular CHO media without any inducers. Each 2-dimensional flow cytometry plot was divided at mCherry=$10^4$ and mCitrine=$10^4$ into four quadrants, representing four states. For each plot, the percentage of cells in each of the four states is labeled on the corresponding corner. Note that in the initial OFF group under the High TMP condition (fourth row, upper set of plots), cell fractions change across time, mainly due to small differences in division rate for cells in different states (data not shown). Timelines above each set of plots represent the indicated inducer conditions.

### Fig. S7

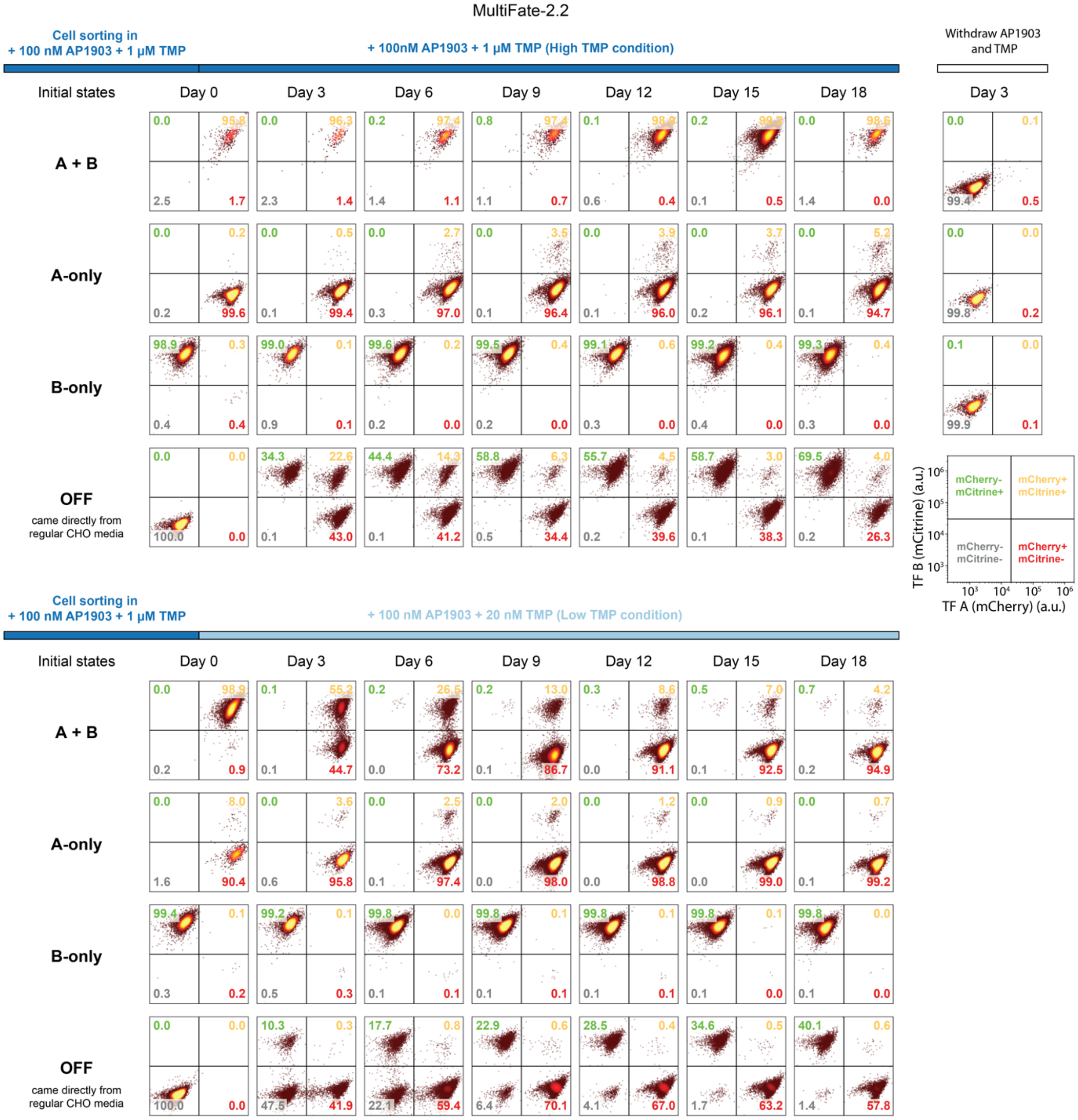
MultiFate-2.2
Cell sorting in
+ 100 nM AP1903 + 1 μM TMP
+ 100nM AP1903 + 1 μM TMP (High TMP condition)
Withdraw AP1903
and TMP
Initial states
Day 0
Day 3
Day 6
Day 9
Day 12
Day 15
Day 18
Day 3
A + B
0.0 95.8 2.5 1.7
0.0 96.3 2.3 1.4
0.2 97.4 1.4 1.1
0.8 97.4 1.1 0.7
0.1 98.9 0.6 0.4
0.2 99.2 0.1 0.5
0.0 98.6 1.4 0.0
0.0 0.1 99.4 0.5
A-only
0.0 0.2 0.2 99.6
0.0 0.5 0.1 99.4
0.0 2.7 0.3 97.0
0.0 3.5 0.1 96.4
0.0 3.9 0.1 96.0
0.0 3.7 0.2 96.1
0.0 5.2 0.1 94.7
0.0 0.0 99.8 0.2
B-only
98.9 0.3 0.4 0.4
99.0 0.1 0.9 0.1
99.6 0.2 0.2 0.0
99.5 0.4 0.2 0.0
99.1 0.6 0.3 0.0
99.2 0.4 0.4 0.0
99.3 0.4 0.3 0.0
0.1 0.0 99.9 0.1
OFF
came directly from
regular CHO media
0.0 0.0 100.0 0.0
34.3 22.6 0.1 43.0
44.4 14.3 0.1 41.2
58.8 6.3 0.5 34.4
55.7 4.5 0.2 39.6
58.7 3.0 0.1 38.3
69.5 4.0 0.2 26.3
mCherry- mCitrine+
mCherry+ mCitrine+
mCherry- mCitrine-
mCherry+ mCitrine-
TF B (mCitrine) (a.u.)
TF A (mCherry) (a.u.)
Cell sorting in
+ 100 nM AP1903 + 1 μM TMP
+ 100 nM AP1903 + 20 nM TMP (Low TMP condition)
Initial states
Day 0
Day 3
Day 6
Day 9
Day 12
Day 15
Day 18
A + B
0.0 98.9 0.2 0.9
0.1 55.2 0.1 44.7
0.2 26.5 0.0 73.2
0.2 13.0 0.1 86.7
0.3 8.6 0.0 91.1
0.5 7.0 0.1 92.5
0.7 4.2 0.2 94.9
A-only
0.0 8.0 1.6 90.4
0.0 3.6 0.6 95.8
0.0 2.5 0.1 97.4
0.0 2.0 0.0 98.0
0.0 1.2 0.0 98.8
0.0 0.9 0.0 99.0
0.0 0.7 0.1 99.2
B-only
99.4 0.1 0.3 0.2
99.2 0.1 0.5 0.3
99.8 0.0 0.1 0.1
99.8 0.1 0.1 0.1
99.8 0.1 0.1 0.1
99.8 0.1 0.1 0.0
99.8 0.0 0.1 0.0
OFF
came directly from
regular CHO media
0.0 0.0 100.0 0.0
10.3 0.3 47.5 41.9
17.7 0.8 22.1 59.4
22.9 0.6 6.4 70.1
28.5 0.4 4.1 67.0
34.6 0.5 1.7 63.2
40.1 0.6 1.4 57.8

**Fig. S7. Raw flow cytometry analysis of the MultiFate-2.2 line.**

Each plot represents one of three replicates at the indicated time point (cf. Fig. 3C). Initial A-only, B-only and A+B cells (rows) were sorted under the media conditions indicated on the top, and initial OFF cells came directly from cells in regular CHO media without any inducers. Each 2-dimensional flow cytometry plot was divided at mCherry=$2x10^4$ and mCitrine=$3x10^4$ into four quadrants, representing four states. For each plot, the percentage of cells in each of the four states is labeled on the corresponding corner. Timelines above each set of plots represent the indicated inducer conditions.

### Fig. S8

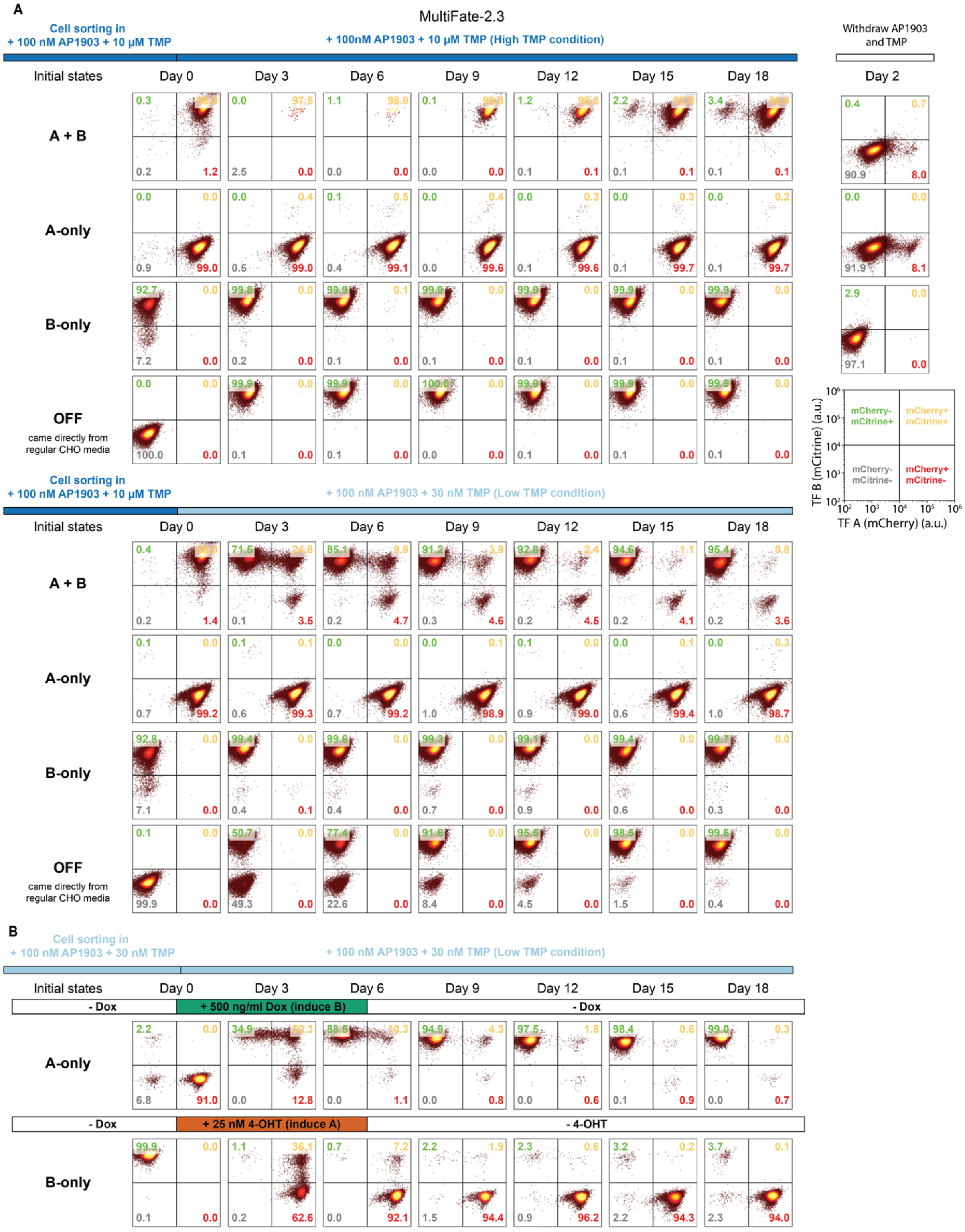
A
MultiFate-2.3
Cell sorting in
+ 100 nM AP1903 + 10 μM TMP
+ 100nM AP1903 + 10 μM TMP (High TMP condition)
Withdraw AP1903 and TMP
Initial states
Day 0
Day 3
Day 6
Day 9
Day 12
Day 15
Day 18
Day 2
A + B
A-only
B-only
OFF
came directly from regular CHO media
mCherry- mCitrine+
mCherry+ mCitrine+
mCherry- mCitrine-
mCherry+ mCitrine-
TF B (mCitrine) (a.u.)
TF A (mCherry) (a.u.)
Cell sorting in
+ 100 nM AP1903 + 10 μM TMP
+ 100 nM AP1903 + 30 nM TMP (Low TMP condition)
B
Cell sorting in
+ 100 nM AP1903 + 30 nM TMP
+ 100 nM AP1903 + 30 nM TMP (Low TMP condition)
- Dox
+ 500 ng/ml Dox (induce B)
- Dox
+ 25 nM 4-OHT (induce A)
- 4-OHT

**Fig. S8. Raw flow cytometry analysis of MultiFate-2.3 stability and switching.**

(A) Each plot represents one of three replicates at the indicated time point (cf. Fig. 3C, F). Initial A-only, B-only and A+B cells (rows) were sorted under the media conditions indicated on the top, and initial OFF cells came directly from cells in regular CHO media without any inducers. Each 2-dimensional flow cytometry plot was divided at mCherry=$10^4$ and mCitrine=$10^4$ into four quadrants, representing four states. For each plot, the percentage of cells in each of the four states is labeled on the corresponding corner. Timelines above each set of plots represent the indicated inducer conditions. (B) To analyze switching, MultiFate-2.3 cells were induced transiently with either 25nM 4-OHT, to induce A expression, or 500 ng/ml Dox (blue timeline), to induce B. 4-OHT or Dox was then washed out and cells were cultured out to t=18 days. Initial A-only, B-only and A+B cells (rows) were sorted under the media conditions indicated on the top, and initial OFF cells came directly from cells in regular CHO media without any inducers.

### Fig. S9

**A The two unstable colonies in MultiFate-2.3 time-lapse images**

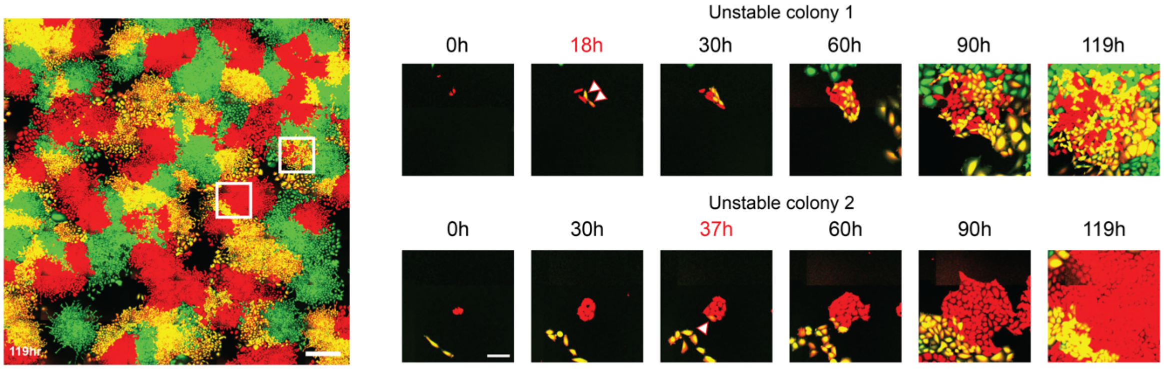


**B The four unstable colonies in MultiFate-3 time-lapse images**

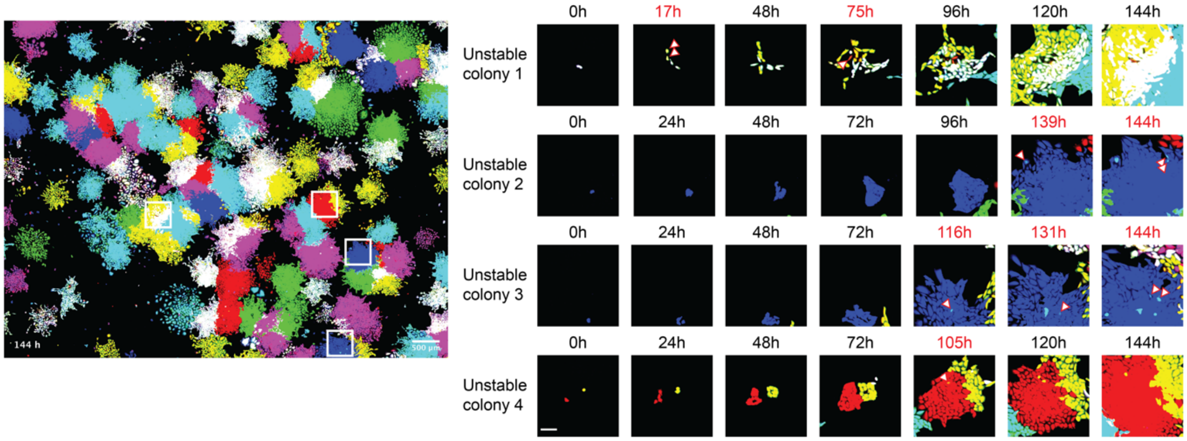

**Fig. S9. Time-lapse movies allow direct visualization of rare spontaneous state-switching events**

(A) Two colonies (white boxes) in a MultiFate-2.3 time-lapse movie exhibited spontaneous state-switching events. Each example is shown as a filmstrip to the right. Arrowheads indicate the cell that switched states. In example 1 (top), a pair of A+B cells (yellow) appear from a colony started in the A-only (red) state (arrowheads). A similar transition was also identified in In example 2 (bottom, white arrowhead). (B) We identified four state-switching events (highlighted in white rectangles) in a MultiFate-3 time-lapse movie. Filmstrips (right) show the events. White arrowheads indicate cells that have switched states. Example 1 (top row) shows a transition from A+B+C (white) to A+B (yellow) and another . Examples 2 and 3 (second row) show a transition from C-only (blue) to B+C (cyan). Example 4 shows a transition from A-only (red) to A+B (yellow). Time points of spontaneous state-switching are labeled in red type. Scale bar is 500 µm for the wide field image (left), and 100 µm for zoomed in images (right).

#### Fig. S10

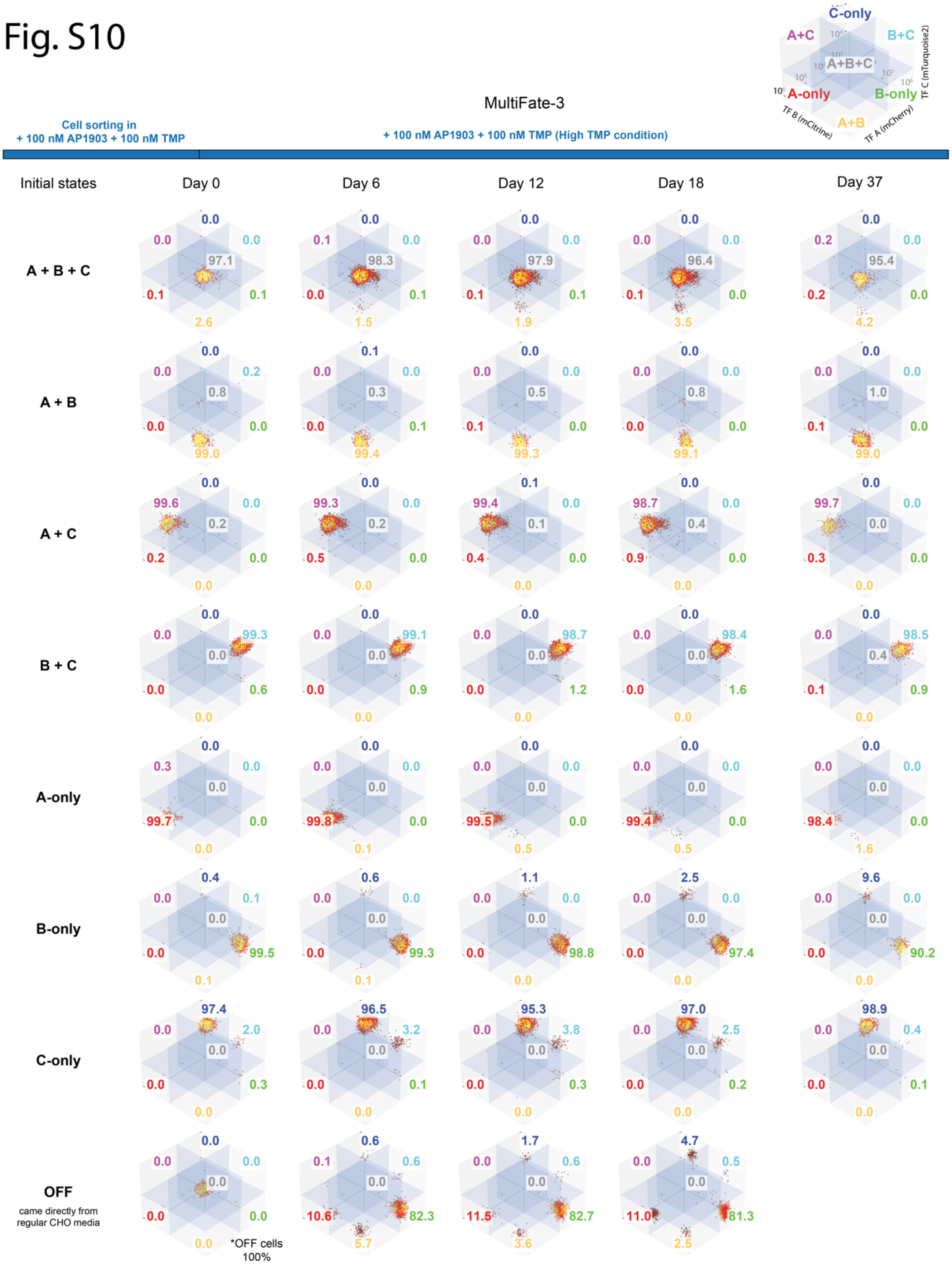
C-only
A+C
B+C
A+B+C
A-only
B-only
A+B
TF C (mTurquoise2)
TF B (mCitrine)
TF A (mCherry)
MultiFate-3
Cell sorting in
+ 100 nM AP1903 + 100 nM TMP
+ 100 nM AP1903 + 100 nM TMP (High TMP condition)
Initial states
Day 0
Day 6
Day 12
Day 18
Day 37
A + B + C
0.0 0.0 0.0 97.1 0.1 0.1 2.6
0.0 0.1 0.0 98.3 0.0 0.1 1.5
0.0 0.0 0.0 97.9 0.1 0.1 1.9
0.0 0.0 0.0 96.4 0.1 0.0 3.5
0.0 0.2 0.0 95.4 0.2 0.0 4.2
A + B
0.0 0.0 0.2 0.8 0.0 0.0 99.0
0.1 0.0 0.0 0.3 0.0 0.1 99.4
0.0 0.0 0.0 0.5 0.1 0.0 99.3
0.0 0.0 0.0 0.8 0.0 0.0 99.1
0.0 0.0 0.0 1.0 0.1 0.0 99.0
A + C
0.0 99.6 0.0 0.2 0.2 0.0 0.0
0.0 99.3 0.0 0.2 0.5 0.0 0.0
0.1 99.4 0.0 0.1 0.4 0.0 0.0
0.0 98.7 0.0 0.4 0.9 0.0 0.0
0.0 99.7 0.0 0.0 0.3 0.0 0.0
B + C
0.0 0.0 99.3 0.0 0.0 0.6 0.0
0.0 0.0 99.1 0.0 0.0 0.9 0.0
0.0 0.0 98.7 0.0 0.0 1.2 0.0
0.0 0.0 98.4 0.0 0.0 1.6 0.0
0.0 0.0 98.5 0.4 0.1 0.9 0.0
A-only
0.0 0.3 0.0 0.0 99.7 0.0 0.0
0.0 0.0 0.0 0.0 99.8 0.0 0.1
0.0 0.0 0.0 0.0 99.5 0.0 0.5
0.0 0.0 0.0 0.0 99.4 0.0 0.5
0.0 0.0 0.0 0.0 98.4 0.0 1.6
B-only
0.4 0.0 0.1 0.0 0.0 99.5 0.1
0.6 0.0 0.0 0.0 0.0 99.3 0.1
1.1 0.0 0.0 0.0 0.0 98.8 0.0
2.5 0.0 0.0 0.0 0.0 97.4 0.0
9.6 0.0 0.0 0.0 0.0 90.2 0.0
C-only
97.4 0.0 2.0 0.0 0.0 0.3 0.0
96.5 0.0 3.2 0.0 0.0 0.1 0.0
95.3 0.0 3.8 0.0 0.0 0.3 0.0
97.0 0.0 2.5 0.0 0.0 0.2 0.0
98.9 0.0 0.4 0.0 0.0 0.1 0.0
OFF
came directly from regular CHO media
0.0 0.0 0.0 0.0 0.0 0.0 0.0
*OFF cells 100%
0.6 0.1 0.6 0.0 10.6 82.3 5.7
1.7 0.0 0.6 0.0 11.5 82.7 3.6
4.7 0.0 0.5 0.0 11.0 81.3 2.5

**Fig. S10. Raw flow cytometry data of MultiFate-3 line under the High TMP condition.**

Each plot represents one of three replicates at the indicated time point (cf. Fig. 4B, High TMP). Initial A-only, B-only and A+B cells (rows) were sorted under the media conditions indicated on the top, and initial OFF cells came directly from cells in regular CHO media without any inducers. Each 3-dimensional flow cytometry plot was divided at mCherry=$2x10^4$, mCitrine=$4x10^4$ and mTurquoise2=$9x10^3$ into eight octants, representing eight states. For each plot, the percentage of cells in each of the 7 states (excluding the OFF state) is labeled on the corresponding octant, as shown in legend at top-right. The timeline (top) represents the indicated inducer conditions. OFF state percentages are usually very low (<1%) across all conditions, and are separately labeled if the percentage is greater than 1%. One of three replicates of cells from each of the 7 initial states (excluding the OFF state) were continuously cultured beyond 18 days. In all 7 states, >90% of cells remained in their original state at day 37 (see indicated percentages).

#### Fig. S11

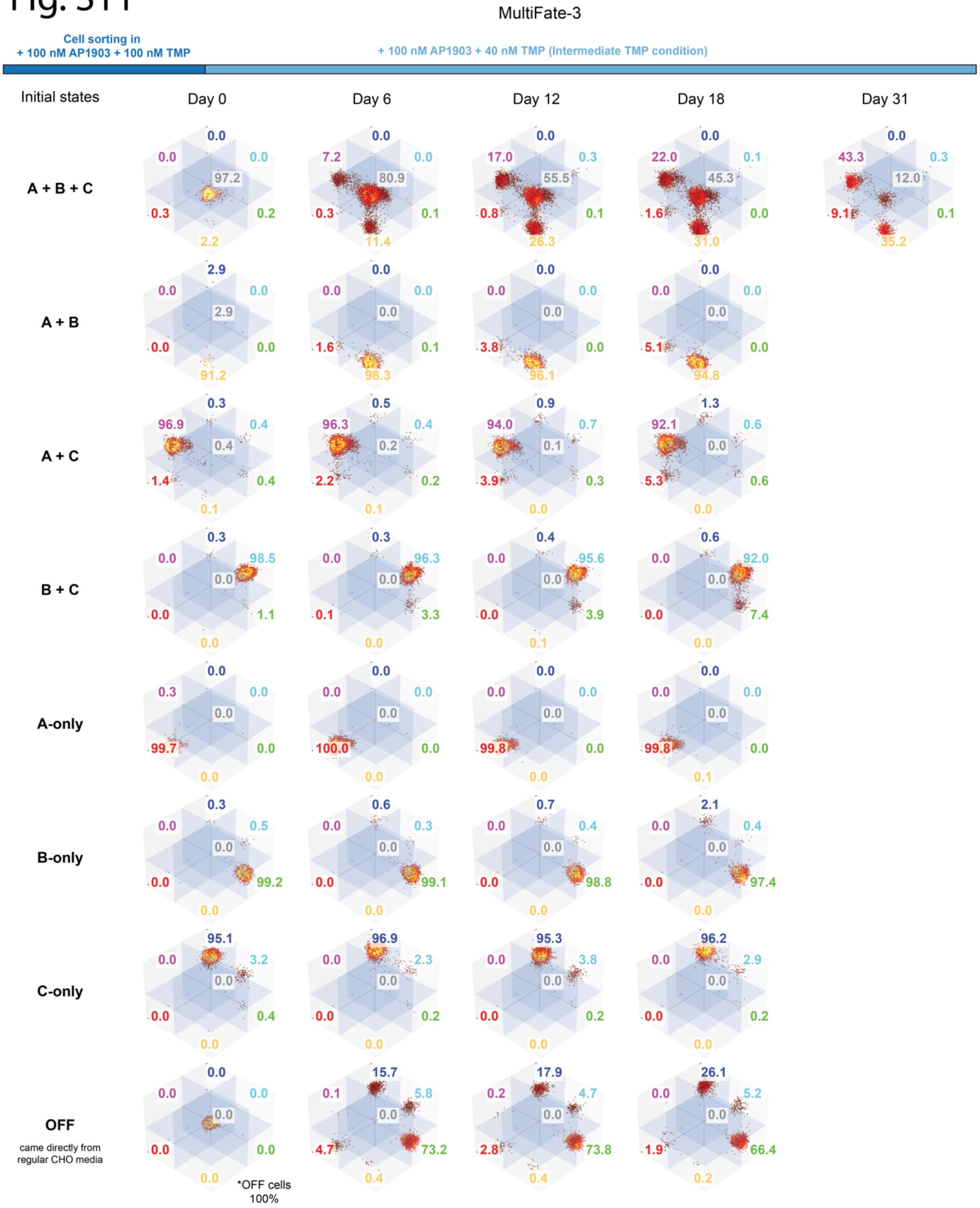
MultiFate-3
Cell sorting in
+ 100 nM AP1903 + 100 nM TMP
+ 100 nM AP1903 + 40 nM TMP (Intermediate TMP condition)
Initial states
Day 0
Day 6
Day 12
Day 18
Day 31
A + B + C
A + B
A + C
B + C
A-only
B-only
C-only
OFF
came directly from regular CHO media
*OFF cells 100%

**Fig. S11. Raw flow cytometry data of MultiFate-3 line under the Intermediate TMP condition.**

Each plot represents one of three replicates at the indicated time point (cf. Fig. 4B, Intermediate TMP). Initial A-only, B-only and A+B cells (rows) were sorted under the media conditions indicated on the top, and initial OFF cells came directly from cells in regular CHO media without any inducers. For each plot, the percentage of cells in each of the 7 states (excluding the OFF state) is labeled on the corresponding octant. The timeline (top) represents the indicated inducer conditions. OFF state percentages are usually very low (<1%) across all conditions, and are separately labeled if the percentage is greater than 1%. Cells from A+B+C initial state were continuously cultured beyond 18 days and measured at day 31. This extended analysis revealed cells escaping from A+B+C state, as predicted, under the Intermediate TMP condition.

### Fig. S12

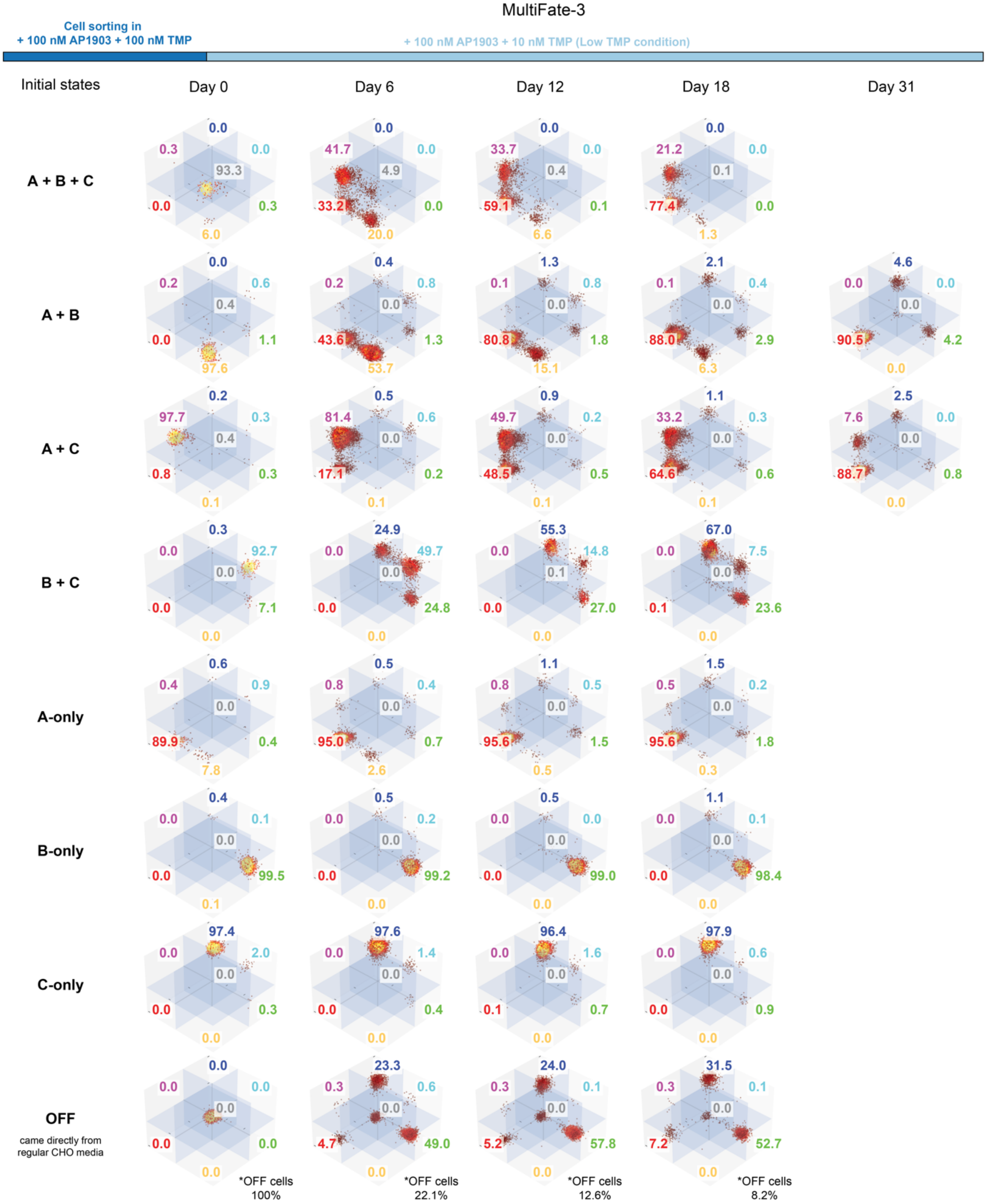

MultiFate-3
Cell sorting in
+ 100 nM AP1903 + 100 nM TMP
+ 100 nM AP1903 + 10 nM TMP (Low TMP condition)
Initial states
Day 0
Day 6
Day 12
Day 18
Day 31
A + B + C
0.0 0.3 0.0 93.3 0.0 0.3 6.0
0.0 41.7 0.0 4.9 33.2 0.0 20.0
0.0 33.7 0.0 0.4 59.1 0.1 6.6
0.0 21.2 0.0 0.1 77.4 0.0 1.3
A + B
0.0 0.2 0.6 0.4 0.0 1.1 97.6
0.4 0.2 0.8 0.0 43.6 1.3 53.7
1.3 0.1 0.8 0.0 80.8 1.8 15.1
2.1 0.1 0.4 0.0 88.0 2.9 6.3
4.6 0.0 0.0 0.0 90.5 4.2 0.0
A + C
0.2 97.7 0.3 0.4 0.8 0.3 0.1
0.5 81.4 0.6 0.0 17.1 0.2 0.1
0.9 49.7 0.2 0.0 48.5 0.5 0.1
1.1 33.2 0.3 0.0 64.6 0.6 0.1
2.5 7.6 0.0 0.0 88.7 0.8 0.0
B + C
0.3 0.0 92.7 0.0 0.0 7.1 0.0
24.9 0.0 49.7 0.0 0.0 24.8 0.0
55.3 0.0 14.8 0.1 0.0 27.0 0.0
67.0 0.0 7.5 0.0 0.1 23.6 0.0
A-only
0.6 0.4 0.9 0.0 89.9 0.4 7.8
0.5 0.8 0.4 0.0 95.0 0.7 2.6
1.1 0.8 0.5 0.0 95.6 1.5 0.5
1.5 0.5 0.2 0.0 95.6 1.8 0.3
B-only
0.4 0.0 0.1 0.0 0.0 99.5 0.1
0.5 0.0 0.2 0.0 0.0 99.2 0.0
0.5 0.0 0.0 0.0 0.0 99.0 0.0
1.1 0.0 0.1 0.0 0.0 98.4 0.0
C-only
97.4 0.0 2.0 0.0 0.0 0.3 0.0
97.6 0.0 1.4 0.0 0.0 0.4 0.0
96.4 0.0 1.6 0.0 0.1 0.7 0.0
97.9 0.0 0.6 0.0 0.0 0.9 0.0
OFF
came directly from regular CHO media
0.0 0.0 0.0 0.0 0.0 0.0 0.0
*OFF cells 100%
23.3 0.3 0.6 0.0 4.7 49.0 0.0
*OFF cells 22.1%
24.0 0.3 0.1 0.0 5.2 57.8 0.0
*OFF cells 12.6%
31.5 0.3 0.1 0.0 7.2 52.7 0.0
*OFF cells 8.2%

**Fig. S12. Raw flow cytometry data of MultiFate-3 line under the Low TMP condition.**

Each plot represents one of three replicates at the indicated time point (cf. Fig. 4B, Intermediate TMP). Initial A-only, B-only and A+B cells (rows) were sorted under the media conditions indicated on the top, and initial OFF cells came directly from cells in regular CHO media without any inducers. For each plot, the percentage of cells in each of the 7 states (excluding the OFF state) is labeled on the corresponding octant. The timeline (top) represents the indicated inducer conditions. OFF state percentages are usually very low (<1%) across all conditions, and are separately labeled if the percentage is greater than 1%. Cells from A+B and A+C initial states were continuously cultured beyond 18 days and measured at day 31. This extended analysis revealed cells escaping from A+B and A+C state, as predicted, under the Low TMP condition.

#### Fig. S13

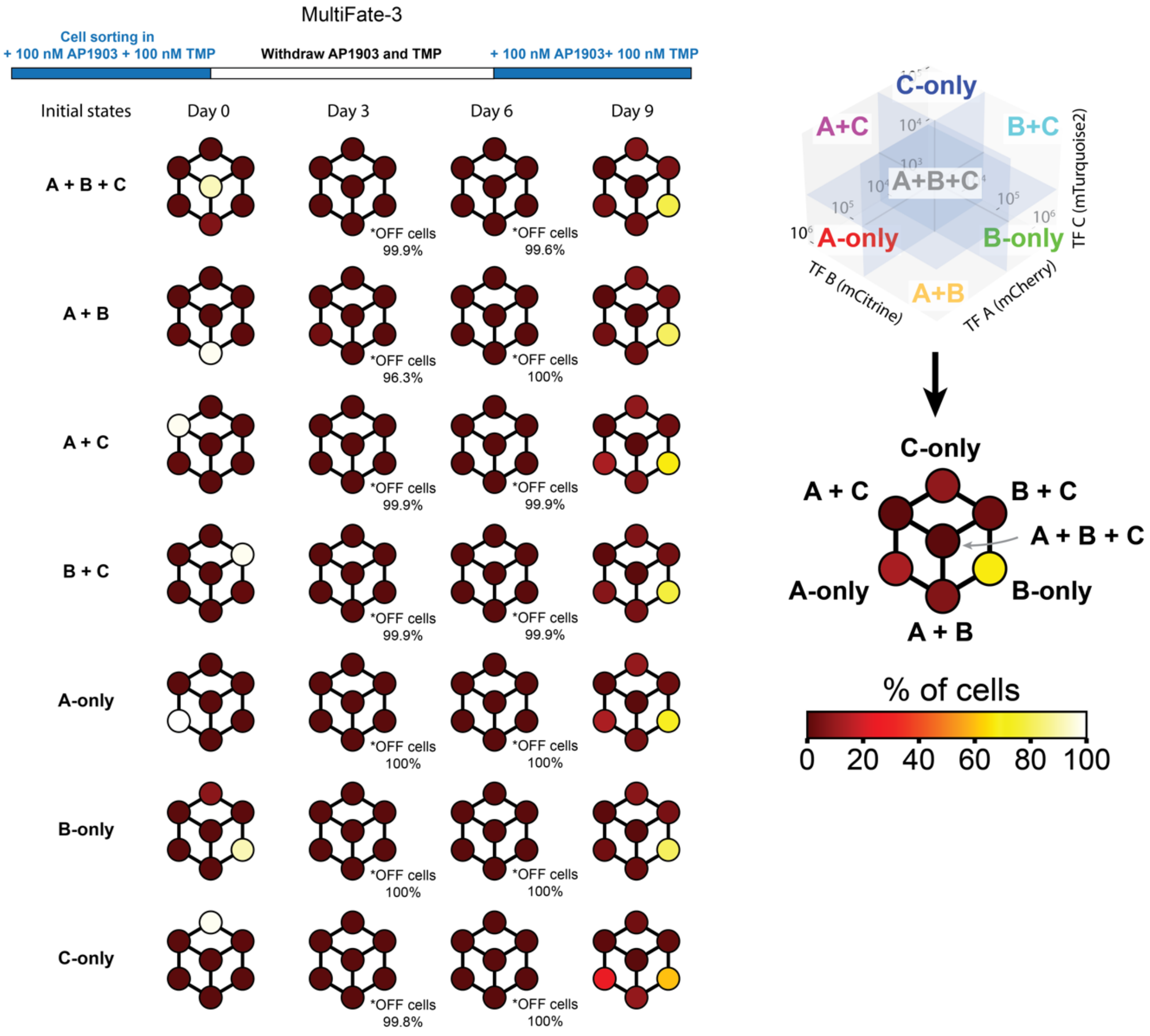
MultiFate-3
Cell sorting in
+ 100 nM AP1903 + 100 nM TMP
Withdraw AP1903 and TMP
+ 100 nM AP1903+ 100 nM TMP
Initial states
Day 0
Day 3
Day 6
Day 9
A + B + C
*OFF cells 99.9%
*OFF cells 99.6%
A + B
*OFF cells 96.3%
*OFF cells 100%
A + C
*OFF cells 99.9%
*OFF cells 99.9%
B + C
*OFF cells 99.9%
*OFF cells 99.9%
A-only
*OFF cells 100%
*OFF cells 100%
B-only
*OFF cells 100%
*OFF cells 100%
C-only
*OFF cells 99.8%
*OFF cells 100%
C-only
A+C
B+C
A+B+C
A-only
B-only
A+B
TF C (mTurquoise2)
TF B (mCitrine)
TF A (mCherry)
C-only
A + C
B + C
A + B + C
A-only
B-only
A + B
% of cells
0 20 40 60 80 100

**Fig. S13. Inducer withdrawal and reintroduction experiments establish dependency on positive autoregulation and rule out the possibility of mixed clones.**

Sorted cells in seven different states were transferred from AP1903+TMP media into regular media without any inducers. Most cells returned to the OFF state within 3 days (second column). After 6 days, AP1903+TMP was added back to the media, and cells were measured by flow cytometry after another 3 days. The resulting state distributions (fourth column) were similar to each other, suggesting that sorted cells in seven different states come from the same monoclonal MultiFate-3 line. Each hexagon represents the mean fractions of three replicates.

Fig. S14

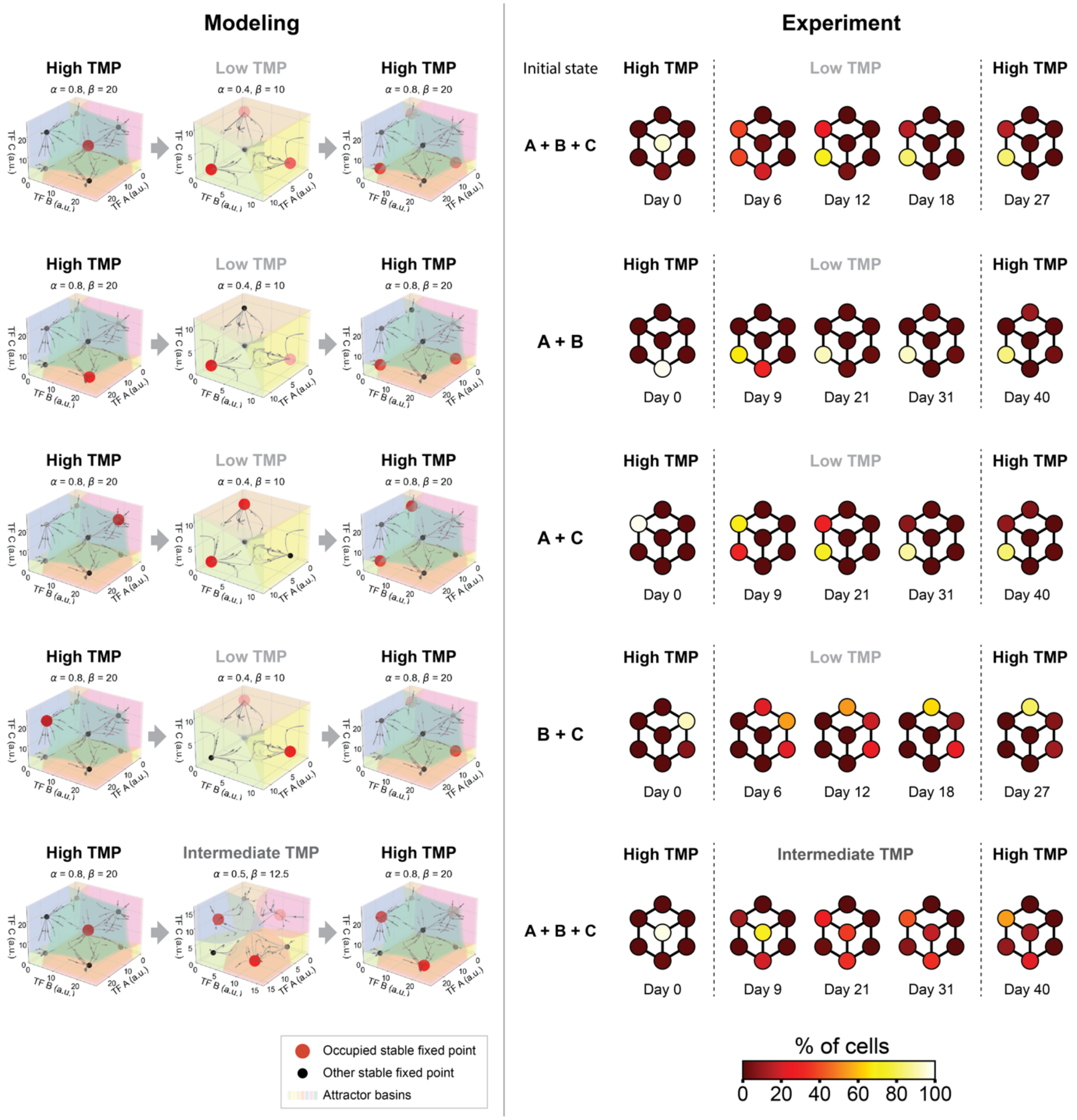

Modeling
Experiment
High TMP
Low TMP
Intermediate TMP
α = 0.8, β = 20
α = 0.4, β = 10
α = 0.5, β = 12.5
TF C (a.u.)
TF B (a.u.)
TF A (a.u.)
Initial state
A + B + C
A + B
A + C
B + C
Day 0
Day 6
Day 9
Day 12
Day 18
Day 21
Day 27
Day 31
Day 40
Occupied stable fixed point
Other stable fixed point
Attractor basins
% of cells
0 20 40 60 80 100

**Fig. S14. MultiFate-3 exhibits predicted hysteresis.**

When transferred from High to Intermediate or Low TMP conditions, cells transition out of destabilized states, as expected. These transitions were irreversible, as shown by both modeling (left) and experiments (right). (Left) Stable fixed points labeled red represent the states that are occupied by simulated cells in the model. The left column shows initial conditions in simulations, with all cells in a single state at High TMP. The middle column shows steady-state density of cells in different states (red intensity) under Low or Intermediate TMP conditions. The right column shows that cells remain in states in the middle column after switching back to the High TMP condition. Phase portraits were calculated using a symmetric MultiFate-3 model (Supplementary Materials). α and β are provided above, $K_d = 1$ and $n = 1.5$. (Right) Experiments showed similar hysteretic behaviors, largely consistent with modeling. In each row, initial cells for indicated states were sorted from the High TMP condition, where they were cultured for at least 3 days, and immediately transferred to Intermediate or Low TMP on day 0. They were then maintained in that condition for 18 or 31 days, as indicated, and then transferred back to the High TMP condition (right). Colorbar indicates density of cells in each of the indicated states, as in Figure 4. Note that a difference between the simulations and experimental results is that actual cells escaping from destabilized states preferentially occupied the A-only state or states containing high A expression, instead of evenly distributing themselves across all states. This reflects some asymmetry of the experimental MultiFate-3 circuit. High TMP condition = 100 nM AP1903 + 100 nM TMP; Intermediate TMP condition = 100 nM AP1903 + 40 nM TMP; Low TMP condition = 100 nM AP1903 + 10 nM TMP. Each hexagon represents the mean fractions of three replicates.

### Fig. S15

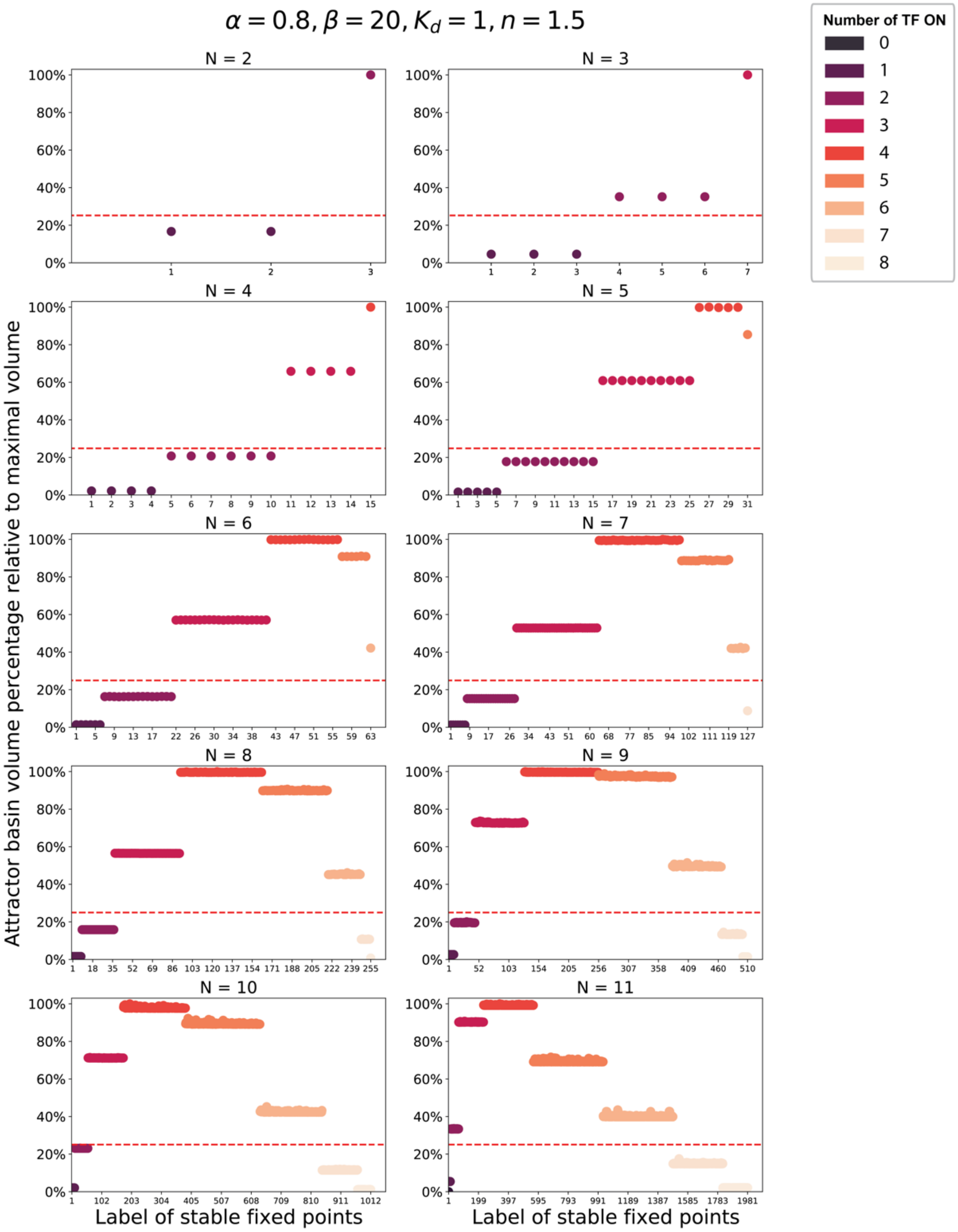

$\alpha = 0.8, \beta = 20, K_d = 1, n = 1.5$
N = 2
N = 3
N = 4
N = 5
N = 6
N = 7
N = 8
N = 9
N = 10
N = 11
Number of TF ON
0
1
2
3
4
5
6
7
8
Attractor basin volume percentage relative to maximal volume
Label of stable fixed points
Label of stable fixed points

**Fig. S15. Additional transcription factors expand the number of accessible stable states.**

Here, we analyzed the number of accessible stable fixed points for systems with different numbers of transcription factors. For each MultiFate-N system, containing N transcription factors, stable fixed points are sorted in ascending order by the number of transcription factors that are ON, which is also indicated by dot colors. For example, in MultiFate-3, the A+B state has 2 transcription factors, A and B, that are ON. Sorted stable fixed points are then labeled from 1, 2, 3, …, k, where k is the total number of stable fixed points for that system. For each stable fixed point, the y-axis value indicates its attractor basin volume (Supplementary Materials) compared with the volume of the biggest attractor basin in the same MultiFate-N system. Stable fixed points that have attractor basin volume at least 25% (red dashed line) of the maximal volume are considered accessible. Since we are using a symmetric MultiFate model, stable fixed points that have the same number of transcription factors ON should have the same attractor basin volumes, resulting in a stair-like pattern in the above plots. Small deviations observable in some cases result from discrete sampling of the phase space.

**Table S1. List of plasmids used in this study and their use in the figures.**

| Index | Construct name | Usage in this study | Figures or MultiFate lines |
|---|---|---|---|
| MF01 | PB-2x(ErbB2bs_ErbB2bs)-TATA-3xNLS-Citrine-BGHpA | Reporter | 2A, S4A,B,C |
| MF02 | PB-2x(37bs_37bs)-TATA-3xNLS-Citrine-BGHpA | Reporter | 2A, S4A,B,C |
| MF03 | PB-2x(42bs_42bs)-TATA-3xNLS-Citrine-BGHpA | Reporter | 2A, S4A,B,C |
| MF04 | PB-2x(92bs_92bs)-TATA-3xNLS-Citrine-BGHpA | Reporter | 2A, S4A,B |
| MF05 | PB-2x(97bs_97bs)-TATA-3xNLS-Citrine-BGHpA | Reporter | 2A, S4A,B |
| MF06 | PB-2x(BCRbs_BCRbs)-TATA-3xNLS-Citrine-BGHpA | Reporter | 2A, S4A,B,C |
| MF07 | PB-2x(HIVbs_HIVbs)-TATA-3xNLS-Citrine-BGHpA | Reporter | 2A, S4A,B |
| MF08 | PB-CAG-ErbB2ZFWT-VP48-mCherry-BGHpA | Trancription factors | 2A, S4A |
| MF09 | PB-CAG-ErbB2ZFWT-GCN4-VP48-mCherry-BGHpA | Trancription factors | 2A, S4A |
| MF10 | PB-CAG-ErbB2ZFR39A-VP48-mCherry-BGHpA | Trancription factors | 2A, S4A |
| MF11 | PB-CAG-ErbB2ZFR39A-GCN4-VP48-mCherry-BGHpA | Trancription factors | 2A, S4A |
| MF12 | PB-CAG-ErbB2ZFR2AR39A-VP48-mCherry-BGHpA | Trancription factors | 2A, S4A |
| MF13 | PB-CAG-ErbB2ZFR2AR39A-GCN4-VP48-mCherry-BGHpA | Trancription factors | 2A, S4A, C |
| MF14 | PB-CAG-ErbB2ZFR2AR39AR67A-VP48-mCherry-BGHpA | Trancription factors | 2A, S4A |
| MF15 | PB-CAG-ErbB2ZFR2AR39AR67A-GCN4-VP48-mCherry-BGHpA | Trancription factors | 2A, S4A |
| MF16 | PB-CAG-FKBP12F36V-BCRZFR39A-VP48-mCherry-BGHpA | Trancription factors | 2B |
| MF17 | PB-CAG-37ZFWT-VP48-mCherry-BGHpA | Trancription factors | S4B |
| MF18 | PB-CAG-37ZFWT-GCN4-VP48-mCherry-BGHpA | Trancription factors | S4B |
| MF19 | PB-CAG-37ZFR39A-VP48-mCherry-BGHpA | Trancription factors | S4B |
| MF20 | PB-CAG-37ZFR39A-GCN4-VP48-mCherry-BGHpA | Trancription factors | S4B |
| MF21 | PB-CAG-37ZFR2AR39A-VP48-mCherry-BGHpA | Trancription factors | S4B |
| MF22 | PB-CAG-37ZFR2AR39A-GCN4-VP48-mCherry-BGHpA | Trancription factors | S4B |
| MF23 | PB-CAG-37ZFR2AR39AR67A-VP48-mCherry-BGHpA | Trancription factors | S4B |
| MF24 | PB-CAG-37ZFR2AR39AR67A-GCN4-VP48-mCherry-BGHpA | Trancription factors | S4B |

| | | | |
|---|---|---|---|
| MF25 | PB-CAG-37ZFR2AR11AR39AR67A-VP48-mCherry-BGHpA | Trancription factors | S4B |
| MF26 | PB-CAG-37ZFR2AR11AR39AR67A-GCN4-VP48-mCherry-BGHpA | Trancription factors | S4B, C |
| MF27 | PB-CAG-42ZFR2AR39AR67A-VP48-mCherry-BGHpA | Trancription factors | S4B |
| MF28 | PB-CAG-42ZFR2AR39AR67A-GCN4-VP48-mCherry-BGHpA | Trancription factors | S4B, C |
| MF29 | PB-CAG-92ZFWT-VP48-mCherry-BGHpA | Trancription factors | S4B |
| MF30 | PB-CAG-92ZFWT-GCN4-VP48-mCherry-BGHpA | Trancription factors | S4B |
| MF31 | PB-CAG-92ZFR39A-VP48-mCherry-BGHpA | Trancription factors | S4B |
| MF32 | PB-CAG-92ZFR39A-GCN4-VP48-mCherry-BGHpA | Trancription factors | S4B |
| MF33 | PB-CAG-92ZFR2AR39A-VP48-mCherry-BGHpA | Trancription factors | S4B |
| MF34 | PB-CAG-92ZFR2AR39A-GCN4-VP48-mCherry-BGHpA | Trancription factors | S4B |
| MF35 | PB-CAG-92ZFR2AR39AR67A-VP48-mCherry-BGHpA | Trancription factors | S4B |
| MF36 | PB-CAG-92ZFR2AR39AR67A-GCN4-VP48-mCherry-BGHpA | Trancription factors | S4B |
| MF37 | PB-CAG-97ZFWT-VP48-mCherry-BGHpA | Trancription factors | S4B |
| MF38 | PB-CAG-97ZFWT-GCN4-VP48-mCherry-BGHpA | Trancription factors | S4B |
| MF39 | PB-CAG-97ZFR39A-VP48-mCherry-BGHpA | Trancription factors | S4B |
| MF40 | PB-CAG-97ZFR39A-GCN4-VP48-mCherry-BGHpA | Trancription factors | S4B |
| MF41 | PB-CAG-97ZFR2AR39A-VP48-mCherry-BGHpA | Trancription factors | S4B |
| MF42 | PB-CAG-97ZFR2AR39A-GCN4-VP48-mCherry-BGHpA | Trancription factors | S4B |
| MF43 | PB-CAG-BCRZF-VP48-mCherry-BGHpA | Trancription factors | S4B |
| MF44 | PB-CAG-BCRZF-GCN4-VP48-mCherry-BGHpA | Trancription factors | S4B |
| MF45 | PB-CAG-BCRZFR39A-VP48-mCherry-BGHpA | Trancription factors | S4B |
| MF46 | PB-CAG-BCRZFR39A-GCN4-VP48-mCherry-BGHpA | Trancription factors | S4B, C |
| MF47 | PB-CAG-HIV1ZFWT-VP48-mCherry-BGHpA | Trancription factors | S4B |
| MF48 | PB-CAG-HIV1ZFWT-GCN4-VP48-mCherry-BGHpA | Trancription factors | S4B |
| MF49 | PB-CAG-HIV1ZFR39A-VP48-mCherry-BGHpA | Trancription factors | S4B |
| MF50 | PB-CAG-HIV1ZFR39A-GCN4-VP48-mCherry-BGHpA | Trancription factors | S4B |
| MF51 | PB-CAG-HIV1ZFR2AR39A-VP48-mCherry-BGHpA | Trancription factors | S4B |

| | | | |
|---|---|---|---|
| MF52 | PB-CAG-HIV1ZFR2AR39A-GCN4-VP48-mCherry-BGHpA | Trancription factors | S4B |
| MF53 | PB-CAG-HIV1ZFR2AR39AR67A-VP48-mCherry-BGHpA | Trancription factors | S4B |
| MF54 | PB-CAG-HIV1ZFR2AR39AR67A-GCN4-VP48-mCherry-BGHpA | Trancription factors | S4B |
| MF55 | PB-CAG-HIV2ZFWT-VP48-mCherry-BGHpA | Trancription factors | S4B |
| MF56 | PB-CAG-HIV2ZFWT-GCN4-VP48-mCherry-BGHpA | Trancription factors | S4B |
| MF57 | PB-CAG-HIV2ZFR39A-VP48-mCherry-BGHpA | Trancription factors | S4B |
| MF58 | PB-CAG-HIV2ZFR39A-GCN4-VP48-mCherry-BGHpA | Trancription factors | S4B |
| MF59 | PB-CAG-HIV2ZFR2AR39A-VP48-mCherry-BGHpA | Trancription factors | S4B |
| MF60 | PB-CAG-HIV2ZFR2AR39A-GCN4-VP48-mCherry-BGHpA | Trancription factors | S4B |
| MF61 | PB-CAG-HIV2ZFR2AR39AR67A-VP48-mCherry-BGHpA | Trancription factors | S4B |
| MF62 | PB-CAG-HIV2ZFR2AR39AR67A-GCN4-VP48-mCherry-BGHpA | Trancription factors | S4B |
| MF63 | PB-TRE3G-6x42bs-6x(BCRbs_BCRbs)-miniCMV-NLS-FKBP12F36V-BCRZFR39A-VP16-NLS-DHFR-IRES-mCitrine-PEST-BGHpA | Self-activation construct | 2C |
| MF64 | PB-TRE3G-6x42bs-6x(37bs_37bs)-miniCMV-NLS-FKBP12F36V-37ZFR2AR11AR39AR67A-VP16-NLS-DHFR-IRES-mCitrine-PEST-BGHpA | Self-activation construct | S5A, MultiFate-2.1, MultiFate-2.3 |
| MF65 | PB-TRE3G-6x42bs-6x(92bs_92bs)-miniCMV-NLS-FKBP12F36V-92ZFR2AR39AR67A-VP16-NLS-DHFR-IRES-mCitrine-PEST-BGHpA | Self-activation construct | S5A |
| MF66 | PB-TRE3G-6x42bs-6x(97bs_97bs)-miniCMV-NLS-FKBP12F36V-97ZFR39A-VP16-NLS-DHFR-IRES-mCitrine-PEST-BGHpA | Self-activation construct | S5A |
| MF67 | PB-TRE3G-6x42bs-6x(ErbB2bs_ErbB2bs)-miniCMV-NLS-FKBP12F36V-ErbB2ZFR2AR39A-VP16-NLS-DHFR-IRES-mCitrine-PEST-BGHpA | Self-activation construct | S5A |
| MF68 | PB-TRE3G-6x42bs-6x(HIVbs_HIVbs)-miniCMV-NLS-FKBP12F36V-HIV1ZFR2AR39A-VP16-NLS-DHFR-IRES-mCitrine-PEST-BGHpA | Self-activation construct | S5A |
| MF69 | PB-TRE3G-6x42bs-6x(HIVbs_HIVbs)-miniCMV-NLS-FKBP12F36V-HIV2ZFR2AR39AR67A-VP16-NLS-DHFR-IRES-mCitrine-PEST-BGHpA | Self-activation construct | S5A |

| | | | |
|---|---|---|---|
| MF70 | PB-TRE3G-6x(42bs_42bs)-miniPromo-42ZFR2AR39AR67A-GCN4-VP48-DHFR-IRES-mCitrine-PEST-BGHpA | Self-activation construct | 2D, S5B |
| MF71 | PB-TRE3G-6x(42bs_42bs)-miniPromo-FKBP12F36V-42ZFR2AR39AR67A-VP48-DHFR-IRES-mCitrine-PEST-BGHpA | Self-activation construct | 2D, S5B |
| MF72 | PB-CAG-IRES-mCherry-PEST-BGHpA (Control) | Protein perturbations | 2D, S5B |
| MF73 | PB-CAG-BCRZFR39A-GCN4-VP48-IRES-mCherry-PEST-BGHpA | Protein perturbations | 2D, S5B |
| MF74 | PB-CAG-FKBP12F36V-BCRZFR39A-VP48-IRES-mCherry-PEST-BGHpA | Protein perturbations | 2D, S5B |
| MF75 | PB-CAG-BCRZFR39A-GCN4-IRES-mCherry-PEST-BGHpA | Protein perturbations | S5B |
| MF76 | PB-CAG-FKBP12F36V-IRES-mCherry-PEST-BGHpA | Protein perturbations | S5B |
| MF77 | PB-CAG-BCRZFR39A-VP48-IRES-mCherry-PEST-BGHpA | Protein perturbations | 2D, S5B |
| MF78 | PB-CAG-BCRZFR39A-IRES-mCherry-PEST-BGHpA | Protein perturbations | S5B |
| MF79 | PB-CAG-GCN4-IRES-mCherry-PEST-BGHpA | Protein perturbations | S5B |
| MF80 | PB-CAG-VP48-IRES-mCherry-PEST-BGHpA | Protein perturbations | S5B |
| MF81 | PB-TRE3G-12x42bs-6x(BCRbs_BCRbs)-miniCMV-NLS-FKBP12F36V-BCRZFR39A-VP16-NLS-DHFR-IRES-mCherry-PEST-BGHpA | Self-activation construct | MultiFate-2.1 |
| MF82 | PB-TRE3G-12x42bs-10x(BCRbs_BCRbs)-miniCMV-NLS-FKBP12F36V-BCRZFR39A-VP16-NLS-DHFR-IRES-mCherry-PEST-BGHpA | Self-activation construct | MultiFate-2.2, MultiFate-3 |
| MF83 | PB-TRE3G-6x42bs-10x(37bs_37bs)-miniCMV-NLS-FKBP12F36V-37ZFR2AR11AR39AR67A-VP16-NLS-DHFR-IRES-mCitrine-PEST-BGHpA | Self-activation construct | MultiFate-2.2, MultiFate-3 |
| MF84 | PB-14xUAS-6x42bs-6x(BCRbs_BCRbs)-miniCMV-NLS-FKBP12F36V-BCRZFR39A-VP16-NLS-DHFR-IRES-mCherry-PEST-BGHpA | Self-activation construct | MultiFate-2.3 |
| MF85 | PB-14xUAS-12x42bs-10x(ErbB2bs_ErbB2bs)-miniCMV-NLS-FKBP12F36V-ErbB2ZFR2AR39A-VP16-NLS-DHFR-IRES-mTurquoise2-PEST-BGHpA | Self-activation construct | MultiFate-3 |
| Note:<br>PB = PiggyBac backbone; TRE3G = Tet3G binding site; UAS = ERT2-Gal4 binding site;<br>TATA, miniCMV, miniPromo are three different minimal promoter;<br>VP48, VP16 are two different transcriptional activation domain;<br>CAG = the constitutive CAG promoter (*87*);<br>NLS = nuclear localization sequence;<br>IRES = internal ribosome entry site; | | | |

BGHpA = bovine growth hormone polyadenylation signal;
PEST = constitutive signal peptide for protein degradation (*114*);
42bs = both the 42ZF binding site and 9bp motif that increase promoter leakiness;
ZFbs_ZFbs = 18bp tandem ZF binding site pairs.

**Table S2. List of stable cell lines constructed for this study and their use in the figures.**

| Cell lines | Parental cells | Poly- or monoclonal | Integrated constructs | Figures | Additional procedures to screen monoclones |
|---|---|---|---|---|---|
| FKBP-BCRZFR39A-VP48-DHFR self-activation | Tet3G-expressing CHO-K1 | Polyclonal | MF63 | 2C | N/A |
| FKBP-37ZFR2AR11AR39AR67A-VP48-DHFR self-activation | Tet3G-expressing CHO-K1 | Polyclonals | MF64 | S5A | N/A |
| FKBP-92ZFR2AR39AR67A-VP48-DHFR self-activation | Tet3G-expressing CHO-K1 | Polyclonal | MF65 | S5A | N/A |
| FKBP-97ZFR39A-VP48-DHFR self-activation | Tet3G-expressing CHO-K1 | Polyclonal | MF66 | S5A | N/A |
| FKBP-ErbB2ZFR2AR39A-VP48-DHFR self-activation | Tet3G-expressing CHO-K1 | Polyclonal | MF67 | S5A | N/A |
| FKBP-HIV1ZFR2AR39A-VP48-DHFR self-activation | Tet3G-expressing CHO-K1 | Polyclonal | MF68 | S5A | N/A |
| FKBP-HIV2ZFR2AR39AR67A-VP48-DHFR self-activation | Tet3G-expressing CHO-K1 | Polyclonal | MF69 | S5A | N/A |
| 42ZFR2AR39AR67A-GCN4-VP48-DHFR self-activation | Tet3G-expressing CHO-K1 | Monoclonal | MF70 | 2D, S5B | Obtained monoclone candidates by limiting dilution, induced candidates with 10 μM TMP and selected the monoclone that spontaneously and homogenously self-activate |
| FKBP-42ZFR2AR39AR67A-VP48-DHFR self-activation | Tet3G-expressing CHO-K1 | Monoclonal | MF71 | 2D, S5B | Obtained monoclone candidates by limiting dilution, induced candidates with 100 nM AP1903 + 10 μM TMP and selected the monoclone that spontaneously and homogenously self-activate |

| | | | | | |
|---|---|---|---|---|---|
| MultiFate-2.1 | ERT2-Gal4-T2A-Tet3G expressing CHO-K1 | Monoclonal | MF64, MF81 | 3C, 3E, S6 | Induced the polyclonal population with 500 ng/ml Dox for 12 hours, then washed out Dox and changed to 100 nM AP1903 + 10 μM TMP for 3 days, FACS sorted monoclones that were mCherry+ and mCitrine+ |
| MultiFate-2.2 | ERT2-Gal4-T2A-Tet3G expressing CHO-K1 | Monoclonal | MF82, MF83 | 3C, S7 | Induced the polyclonal population with 500 ng/ml Dox for 12 hours, then washed out Dox and changed to 100 nM AP1903 + 10 μM TMP for 3 days, FACS sorted monoclones that were mCherry+ and mCitrine+ |
| MultiFate-2.3 | ERT2-Gal4-T2A-Tet3G expressing CHO-K1 | Monoclonal | MF64, MF84 | 3C, 3D, 3F, S8 | Induced the polyclonal population with 500 ng/ml Dox and 75 nM 4-OHT for 12 hours, then washed out Dox and 4-OHT and changed to 100 nM AP1903 + 10 μM TMP for 3 days, FACS sorted monoclones that were mCherry+ and mCitrine+ |
| MultiFate-3 | MultiFate-2.2 | Monoclonal | MF85 | 4B, 4C, S10-S14 | Induced the polyclonal population with 500 ng/ml Dox and 75 nM 4-OHT for 12 hours, then washed out Dox and 4-OHT and changed to 100 nM AP1903 + 10 μM TMP for 3 days, FACS sorted monoclones that were mCherry+, mCitrine+ and mTurquoise2+ |
| Promoter structures of different MultiFate lines:<br>MultiFate-2.1<br>TF A promoter has Tet3G binding sites, 6x(BCRbs_BCRbs);<br>TF B promoter has Tet3G binding sites, 6x(37bs_37bs);<br>MultiFate-2.2<br>TF A promoter has Tet3G binding sites, 10x(BCRbs_BCRbs);<br>TF B promoter has Tet3G binding sites, 10x(37bs_37bs);<br>MultiFate-2.3<br>TF A promoter has ERT2-Gal4 binding sites, 6x(BCRbs_BCRbs);<br>TF B promoter has Tet3G binding sites, 6x(37bs_37bs);<br>MultiFate-3<br>TF A promoter has Tet3G binding sites, 10x(BCRbs_BCRbs);<br>TF B promoter has Tet3G binding sites, 10x(37bs_37bs);<br>TF C promoter has ERT2-Gal4 binding sites, 10x(ErbB2bs_ErbB2bs); | | | | | |

**Table S3. List of molecular reactions and their propensities for Gillespie simulation.**

| Reactions | Molecule update | Propensity |
|---|---|---|
| TF A mRNA transcription | $a \rightarrow a + 1$ | $k_1 + k_2 \frac{[A_2]^n}{K_M^n + [A_2]^n}$ |
| TF A mRNA removal | $a \rightarrow a - 1$ | $\delta_{mRNA}[a]$ |
| TF B mRNA transcription | $b \rightarrow b + 1$ | $\mathrm{r}k_1 + \mathrm{m}k_2 \frac{[B_2]^n}{(\kappa \mathrm{K}_M)^n + [B_2]^n}$ |
| TF B mRNA removal | $b \rightarrow b - 1$ | $\delta_{mRNA}[b]$ |
| TF A protein translation | $A \rightarrow A + 1$ | $k_p[a]$ |
| TF A protein removal | $A \rightarrow A - 1$ | $\delta[A]$ |
| TF B protein translation | $B \rightarrow B + 1$ | $k_p[b]$ |
| TF B protein removal | $B \rightarrow B - 1$ | $\gamma\delta[B]$ |
| TF A homodimerization | $A \rightarrow A - 2, A_2 \rightarrow A_2 + 1$ | $k_{on}[A]^2 \times ([A] \geq 2)$ |
| AA homodimer dissociation | $A_2 \rightarrow A_2 - 1, A \rightarrow A + 2$ | $k_{off}[A_2]$ |
| TF B homodimerization | $B \rightarrow B - 2, B_2 \rightarrow B_2 + 1$ | $k_{on}[B]^2 \times ([B] \geq 2)$ |
| BB homodimer dissociation | $B_2 \rightarrow B_2 - 1, B \rightarrow B + 2$ | $k_{off}[B_2]$ |
| Heterodimerization | $A \rightarrow A - 1, B \rightarrow B - 1, AB \rightarrow AB + 1$ | $2k_{on}[A][B]$ |
| AB heterodimer dissociation | $AB \rightarrow AB - 1, A \rightarrow A + 1, B \rightarrow B + 1$ | $k_{off}[AB]$ |
| AA homodimer removal | $A_2 \rightarrow A_2 - 1$ | $\delta[A_2]$ |
| BB homodimer removal | $B_2 \rightarrow B_2 - 1$ | $\gamma\delta[B_2]$ |
| AB –> B due to A removal | $AB \rightarrow AB - 1, B \rightarrow B + 1$ | $\delta[AB]$ |
| AB –> A due to B removal | $AB \rightarrow AB - 1, A \rightarrow A + 1$ | $\gamma\delta[AB]$ |

Note:
A, B, $A_2$, $B_2$, AB represents proteins of monomer A, monomer B, homodimer AA, homodimer BB and heterodimer AB, respectively.
a, b represents mRNAs of A and B, respectively.

**Table S4. List of physiologically reasonable parameter regimes.**

| Parameters | Model | Estimated values | References |
|---|---|---|---|
| $K_M$ | Deterministic and stochastic | 10 nM or 8000 molecules/cell | (*91–94*) |
| δ | Deterministic and stochastic | 0.1 $hr^{-1}$ for "High TMP" condition; 0.2 $hr^{-1}$ for "Low TMP" condition | (*61*, *95*, *96*) |
| $n$ | Deterministic and stochastic | 1.5 | (*91*, *101–103*, *105*, *115*) |
| α | Deterministic | 0.8 nM/hr | Derived |
| β | Deterministic | 20 nM/hr | Derived |
| $K_d$ | Deterministic | 10 nM | (*106*) and current study |
| $k_1$ | Stochastic | 3.2 mRNA/hr | Current study |
| $k_2$ | Stochastic | 80 mRNA/hr | (*116*) and current study |
| $\delta_{mRNA}$ | Stochastic | 0.7 $hr^{-1}$ | (*97*, *99*, *100*) |
| $k_p$ | Stochastic | 140 proteins/(mRNA*hr) | (*97*) |
| $k_{on}$ | Stochastic | 1.8x$10^{-3}$ cell/(protein*hr) | (*107*) |
| $k_{off}$ | Stochastic | 14.4 $hr^{-1}$ | Derived |
| Note: 1 nM is equivalent to 800 molecules per CHO cell. | | | |

**Movie S1. Modeling of MultiFate-2 state-switching dynamics.**

This simulation shows the phase plane and dynamics of individual cells (red dots) for indicated parameters (top) in the non-dimensionalized MultiFate-2 model (Box 1). The movie shows a symmetric regime, switching transiently to an asymmetric regime and then returning to the symmetric regime once again. Parameter r (Supplementary Materials) represents the ratio of basal protein production rate between TF A and TF B (r>1 means higher B basal production). r= 1 at the beginning and end of the movie, and r= 5 in the middle frames. Frame interval is 2 time units in the Gillespie algorithm (Material and Methods). "B overexpression" refers to elevate basal expression of B (r>1). Selected frames are presented in Fig. S3.

**Movie S2. Time-lapse movie of MultiFate-2.3 cells in 3 stable states.**

MultiFate-2.3 cells from A-only (mCherry, shown in red), B-only (mCitrine, shown in green) and A+B (mCherry+mCitrine, shown in yellow) states were pre-mixed at 1:1:1 ratio. 2000 cells from the cell mixture were plated sparsely in one well of a 24-well plate with media containing 100 nM AP1903 and 10 μM TMP. The movie started 12 hours after cell plating and lasted for 5 days. Selected frames from this movie are presented in Fig. 3D.

**Movie S3. Modeling of MultiFate-2 bifurcation dynamics and irreversible transitions.**

Simulation of MultiFate-2 model (Box 1) for symmetric (left) and asymmetric (right) parameter sets. In both cases, the system transitions from a tristable regime to a bistable regime, and then returns to the initial tristable regime. Red dots represent individual stochastically simulated cells. Irreversible transitions in both cases, but produce asymmetric cell state distributions in the asymmetric case (right). Frame interval is 2 time units in the Gillespie algorithm (Material and Methods). Non-dimensionalized parameter sets chosen for the High TMP condition and the Low TMP condition are the same as those used in the main figures. Parameter is one of asymmetric parameters (Supplementary Materials), which represent the ratio of TF concentrations for half-maximal activation between TF A and TF B (>1 indicates that higher B concentrations needed for half-maximal activation, B is a weaker activator). Modeling shows that cells are preferentially attracted to the A-only state in the asymmetric MultiFate-2 circuit during bifurcation, explaining why experimental MultiFate-2 cells are unevenly distributed between A-only and B-only states after escaping from destabilized A+B state under the Low TMP condition. Selected frames for symmetric MultiFate-2 are presented in Fig. 3E.

**Movie S4. MultiFate-3 Phase Diagrams.**

These phase diagrams, which are the same as in Figs. 1D and S2A, are slowly rotated. All phase portraits were generated using the symmetric MultiFate-3 model with non-dimensionalized parameters.

**Movie S5. Time-lapse movie of MultiFate-3 cells in 7 stable states.**

MultiFate-3 cells from A-only (mCherry, shown in red), B-only (mCitrine, shown in green), C-only (mTurquoise2, shown in blue), A+B (yellow), A+C (magenta), B+C (cyan) and A+B+C (white) states were pre-mixed at equal ratio. 800 cells from cell mixture were plated in one well of a 24-well plate with media containing 100 nM AP1903 and 100 nM TMP. The movie started 6 hours after cell plating and lasted for 6 days. Selected frames are presented in Fig. 4C.
